# The histidine kinase PdtaS is a cyclic di-GMP binding metabolic sensor that controls mycobacterial adaptation to nutrient deprivation

**DOI:** 10.1101/615575

**Authors:** Vignesh Narayan Hariharan, Chandrani Thakur, Albel Singh, Renu Gopinathan, Devendra Pratap Singh, Gaurav Sankhe, Nagasuma Chandra, Apoorva Bhatt, Deepak Kumar Saini

## Abstract

Cell signalling relies on second messengers to transduce signals from the sensory apparatus to downstream components of the signalling pathway. In bacteria, one of the most important and ubiquitous second messengers is the small molecule cyclic diguanosine monophosphate (c-di-GMP). While the biosynthesis, degradation and regulatory pathways controlled by c-di-GMP are well characterized, the mechanisms through which c-di-GMP controls these processes is not completely understood. Here we present the first report of a c-di-GMP regulated sensor histidine kinase previously named PdtaS (Rv3220c), which binds to c-di-GMP at sub-micromolar concentrations, subsequently perturbing signalling of the PdtaS-PdtaR (Rv1626) two component system. Aided by biochemical analysis, molecular docking and structural modelling, we have characterized the binding site of c-di-GMP in the GAF domain of PdtaS. We show that a *pdtaS* knockout in *M. smegmatis* is severely compromised in growth on amino acid deficient media and exhibits global transcriptional dysregulation. Perturbation of the c-di-GMP-PdtaS-PdtaR axis results in a cascade of cellular changes recorded by a multi-parametric systems approach of transcriptomics, unbiased metabolomics and lipid analyses.

**One-sentence summary:** The universal bacterial second messenger cyclic di-GMP controls the mycobacterial nutrient stress response

## Introduction

*Mycobacterium tuberculosis* (Mtb), the causative agent of a disease that is now the leading cause of death due to a single infectious agent, is estimated to have infected 10.4 million people worldwide in 2016 (1). Two component signalling systems (TCSs) play an important role in multiple stages of the Mtb lifestyle within the host including regulating viability, host cell invasion, intracellular survival, phago-lysosomal fusion, dormancy and cell division (2). Moreover, the complete absence of TCSs in mammals and their importance in Mtb physiology also makes them attractive targets for anti-tubercular therapy (3, 4).

However, the functional study of TCS signal transduction is confounded by a lack of information regarding ligands that activate TCS sensor histidine kinases (SKs). While broad cues that activate TCSs are known, like the pH responsive PhoR-PhoP TCS or the iron responsive TcrY-TcrX TCS (5), ten out of thirteen SKs in Mtb have no known ligands and therefore no way to specifically activate the TCS to study downstream signalling. Addressing this lacuna in our understanding of SKs will go a long way towards delineating the importance of TCSs in Mtb physiology and the rational design of inhibitors against TCS signalling.

Among the less understood TCSs of Mtb is the highly conserved PdtaS-PdtaR TCS. Studies have shown PdtaR to be essential for survival on medium containing cholesterol as the sole carbon source (6) and intracellular growth during macrophage infection (7). Its cognate SK PdtaS has been implicated in controlling ribosomal protein composition and sensitivity to ribosome targeting antibiotics, membrane transport inhibitors and respiratory chain antagonists in *Mycobacterium smegmatis* (8). However, neither the ligands that triggers the activation of this TCS nor the adaptive responses that result from PdtaS-PdtaR signalling are known.

In this study, we use a knockout of *pdtaS* in *Mycobacterium smegmatis* to identify the role of this conserved TCS in mycobacterial physiology. Using comparative metabolomics and in silico docking, we have identified c-di-GMP as a ligand for PdtaS. Furthermore, we show that PdtaS phosphorylation and integrity of the c-di-GMP binding site are essential for growth, lipid homeostasis and transcriptional adaptation during nutrient deprivation. Finally, using protein-protein interaction network modelling of the transcriptomic data, we uncover a novel interactome that connects diverse metabolic pathways in mycobacteria.

## Results

### PdtaS deficiency results in poor growth during nutrient deprivation

In order to study the role of PdtaS and PdtaS-PdtaR signalling in the physiology of mycobacteria, we deleted *pdtaS* (MSMEG_1918) in *M. smegmatis* using specialized transduction as described previously (9) and verified the deletion using PCR and Southern blot analysis (Fig. 1A and Fig. S1). In addition, the *M. smegmatis pdtaS* knockout strain (Δms*pdtaS*), was complemented with *M. tuberculosis pdtaS* (mt*pdtaS*) driven by its native promoter using the integrative vector pCV125 (Δ:mt*pdtaS*) to study the functions of mtPdtaS.

**Figure 1.**
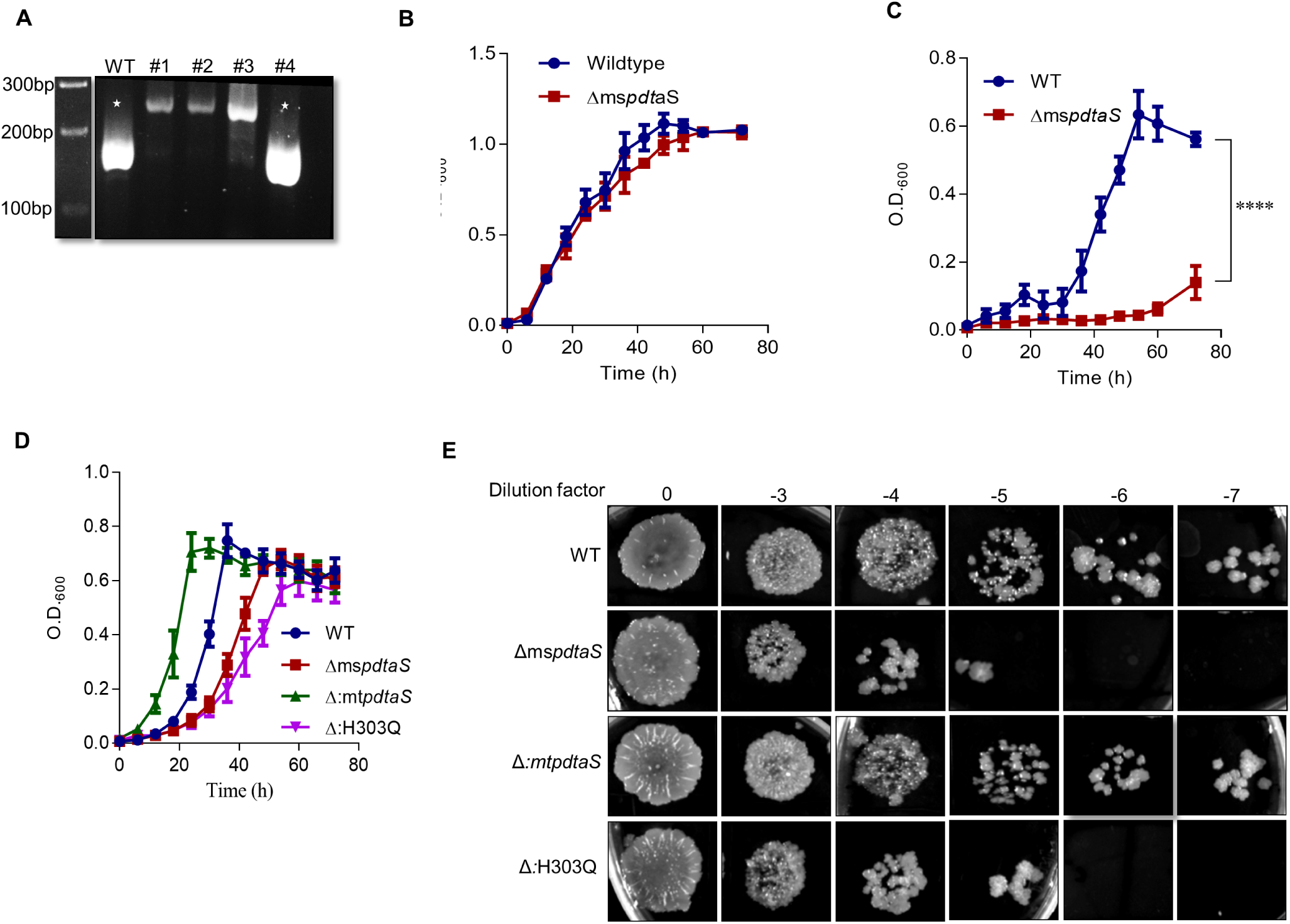
PdtaS activity is essential for growth during nutrient deprivation. **a.** Generation of PdtaS knockout *M. smegmatis* strain. PCR analysis of genomic DNA isolated from wildtype (WT) and *pdtaS* knockout colonies (#1-#4) for verification of gene knockouts. White stars indicate amplification product of wildtype loci. Southern blot analysis for the same can be found in Figure S1. **b, c, d**. Growth curves of various cultures. The optical density at 600nm (O.D._600_) was measured at specific time points for wildtype (WT), *pdtaS* knockout (Δms*pdtaS*), complemented (Δ:mt*pdtaS*) and phosphorylation-defective (Δ:H303Q) strains grown in **b.** Tryptic soy broth, **c.** 7H9 + 0.2% glycerol **d.** 7H9 + 0.2% glycerol after pre-inoculation in tryptic soy broth. Error bars indicate the standard error of the mean of biological triplicates. Identification and validation of phosphorylation sites can be found in Figure S2. Growth curves after starvation of pre-inoculum can be found in Figure S3. Statistical significance as measured by unpaired t-tests for each row is depicted as ns – not significant, **** – p<0.0001 **e.** Dilution spot assay. Strains grown in 7H9 + 0.2% glycerol after pre-inoculation in tryptic soy broth were diluted and spotted onto agar plates to assess the viable colony forming units.

To understand the role of signalling in mtPdtaS function, the conserved histidine residue involved in signal transduction was identified and verified biochemically by radiolabelled kinase assays to be His^303^ (Fig. S2). This residue was mutated to Gln in the copy of mt*pdtaS* on the complementation plasmid and electroporated into Δms*pdtaS* (Δ:H303Q) to study the importance of phosphorylation in mtPdtaS function. The *M. smegmatis pdtaS* knockout strain did not show any significant difference in growth compared to the wildtype strain when grown in the nutrient rich tryptic soy broth (TSB) medium (Fig. 1B), similar to a previous report (8). However, when single colonies of Δms*pdtaS* were grown in nutritionally poorer 7H9 broth base containing only glycerol, very little growth was observed even during prolonged incubation periods (Fig. 1C). In order to investigate if the poor growth in 7H9 + glycerol (henceforth called poor medium) was related to the availability of nutrients, strains were allowed to grow till stationary phase in TSB to ensure high levels of intracellular nutrients and then sub-cultured into poor medium. It was found that the severity of the growth defect of Δms*pdtaS* was ameliorated when cultures were pre-grown in TSB and then shifted to poor medium containing glycerol as the sole carbon source. Complementation of the knockout with the wildtype *M. tuberculosis pdtaS* (Δ:mt*pdtaS*) was able to rescue the growth but complementation with biochemically inactive mt*pdtaS* (Δ:H303Q) failed to do so (Fig. 1D). The growth defect was also captured by dilution spotting analysis at 36h of growth in poor medium, with WT colony forming units (cfu) two logs greater than those of Δms*pdtaS* (Fig. 1E). Additionally, when stationary phase TSB grown cultures were carbon starved for 8h (resuspended in 7H9 alone) to deplete intracellular nutrient pools and then sub-cultured in poor medium containing either glucose or glycerol as the sole carbon source, a larger difference between the growth rates of WT and Δms*pdtaS* (Fig. S3) was recorded, indicating that the growth defect directly correlated with intracellular nutrient pools.

### PdtaS signalling remodels the global transcriptome of *M. smegmatis*

The activation of PdtaS leads to phosphorylation of its cognate response regulator (RR), PdtaR (10, 11) which presumably binds to target RNA through its RNA binding ANTAR domain to effect transcription anti-termination of target genes (12). Previous reports have shown that mtPdtaS does not phosphorylate any other RR, making it a highly specific TCS (11). In order to study the processes affected by the absence of PdtaS and therefore phosphorylated PdtaR, a gene expression microarray analysis was performed for WT and Δms*pdtaS* strains grown in 7H9 containing glucose as the sole carbon source (Fig. 2A). A list of statistically significant Differentially Expressed Genes (DEGs) can be found in Supplementary File 1.

**Figure 2.**
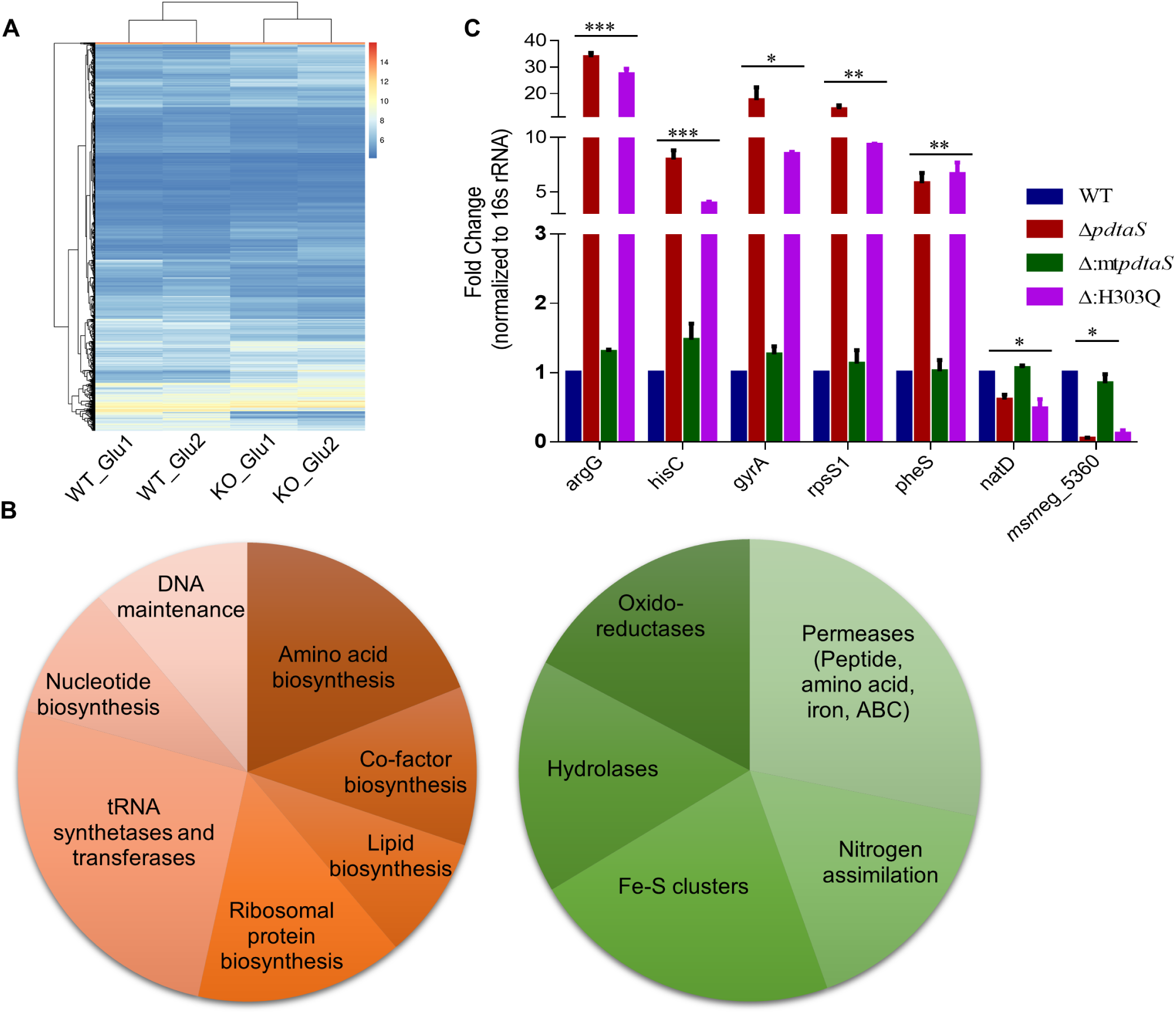
Absence of active PdtaS results in global transcriptomic dysregulation. **a.** Heatmap showing hierarchical clustering of log2 normalized gene expression values of microarray data performed in biological duplicates. Gene expression levels in wild type strain are marked as WT and KO indicates *pdtaS* knockout strain. **b.** Pie-chart of differentially regulated processes. Upregulated processes are shaded red (left) and downregulated processes are shaded green (right). **c.** qRT-PCR analysis of genes representative of microarray DEGs. Relative expression levels of *argG, hisC, gyrA, rpsS1, pheS, natD* and *msmeg*_5360 in RNA isolated from wildtype *M. smegmatis*, Δms*pdtaS* mutant and the complemented strains Δ*pdtaS*:mt*pdtaS* and Δ:H303Q is depicted. Expression is normalized to 16s rRNA gene expression as an internal control. The fold change in gene expression in the Δms*pdtaS* mutant relative to wildtype is plotted as mean of three independent experiments. Error bars represent standard error. The expression of genes in the wildtype is set to 1. *** – p<0.001 ** – p<0.01* – p<0.05.

Significant DEGs were manually grouped into gene families to identify processes that were affected by the absence of PdtaS. The main processes downregulated in the Δms*pdtaS* strain were the Fe-S cluster family, hydrolases, oxido-reductases, transmembrane transport and nitrogen assimilation genes. Genes upregulated belonged to the processes of nucleotide biosynthesis, DNA maintenance, amino acid biosynthesis, co-factor biosynthesis, fatty acid biosynthesis, ribosomal protein biosynthesis and tRNA synthetases and transferases (Fig. 2B). The analyses revealed that the Δms*pdtaS* strain appeared to deficient in adaptations that are required for growth in nutrient poor conditions such as the downregulation of macromolecular biosynthesis and translation with a concomitant upregulation in nutrient uptake and macromolecular salvage enzymes such as hydrolases.

Selected differentially regulated genes from the microarray data were validated using quantitative reverse transcriptase PCR. It was found that amino acid biosynthesis, represented by members of the arginine (*argG*) and histidine (*hisC*) biosynthesis operons, DNA gyrase (*gyrA*) important for replication and transcription and small ribosomal protein *rpsS1*, phenylalanylyl tRNA synthetase *pheS* and the initiator tRNA, tRNA-fMet, representative of the translation machinery were all upregulated five-folds or more in Δms*pdtaS* and Δ:H303Q compared to WT and Δ:mt*pdtaS* (Fig. 2c).

### PdtaS deficiency results in a reduction of FASI products

The outcome of transcriptional dysregulation on the physiology of Δms*pdtaS* cells was studied using a comparative metabolomics approach using apolar and polar fractions of the intracellular metabolome. Interestingly, despite the large number of processes affected at the transcriptional level, the metabolomic differences between WT and Δms*pdtaS* strains were principally in two classes of metabolites (Supplementary file 2). Details of all metabolites identified in the study can be found in supplementary file 3. The greatest change in metabolome was the reduction in levels of multiple lipid classes in the Δms*pdtaS* strain. Lipids identified by the metabolomics were categorized into the following classes – acyl glycerides, fatty acids, alcohols and esters, glycerophospholipids and sialylated glycosphingolipids (details in Supplementary file 2). Metabolites that belong to more than one lipid class were categorized separately as mixed lipid classes. The levels of metabolites in four out of these five lipid classes were greatly reduced (Table 1 and Fig. 3A). Individual species of two lipid classes, acyl glycerides and phospholipids have been plotted as representative of the large-scale reduction in lipids in Δms*pdtaS* compared to WT (Fig. 3B and 3C). Second, the soluble fraction of Δms*pdtaS* was enriched in dipeptides (Fig. 3D), indicating higher levels of protein degradation or reduction in dipeptide exporters in the *pdtaS* knockout. A previous study has demonstrated dipeptide accumulation to have a growth inhibitory effect in *E. coli* (13), and could therefore be contributing to the poor growth of Δms*pdtaS* cells.

**Figure 3.**
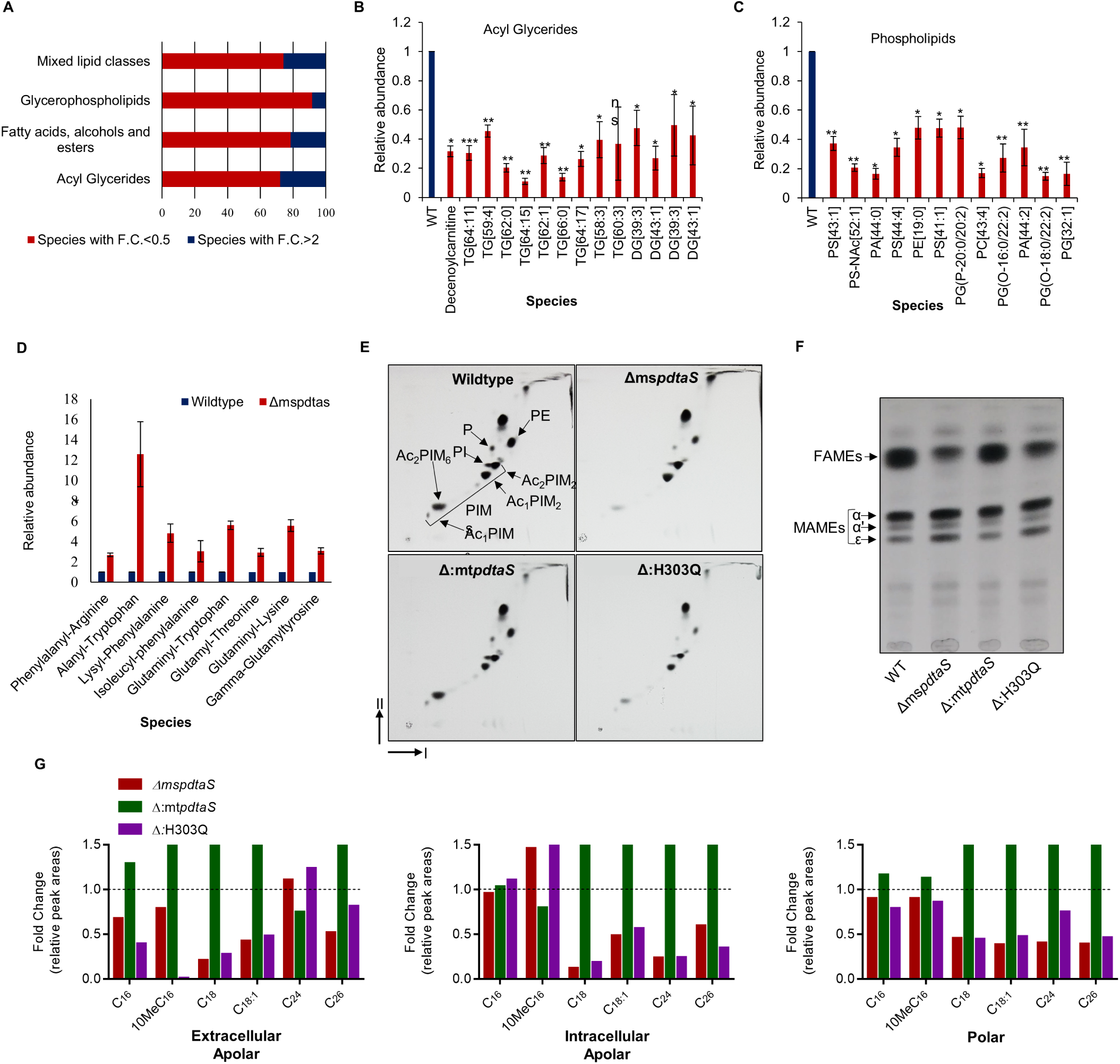
Loss of PdtaS signalling results in dysregulation of lipid and protein homeostasis. **a**. Abundance ratios of lipid classes. The percentage of metabolites that are replete or depleted in Δ*ms*pdtaS cells compared to wildtype *M. smegmatis* in each lipid class. Fold change (F.C.) is calculated as relative quantity of each metabolite in Δms*pdtaS* cells with respect to wildtype cells. Only species with p<0.05 have been represented. **b, c**. The abundance of individual metabolites categorized as **b**. acyl glycerides and **c**. phospholipids in Δms*pdtaS* cells relative to wildtype cells. WT-wildtype, TG-triacylglyceride, DG-diacylglyceride, PS-phosphatidyl serine, PA-phosphatidic acid, PE-phosphatidyl ethanolamine, PG-phosphatidyl glyceride. *** – p<0.001 ** – p<0.01 * – p<0.05 ns – not significant. **d**. Relative abundance of dipeptides. The abundance of individual dipeptides in Δms*pdtaS* cells relative to wildtype cells. **e**. 2D-TLC analysis of polar lipids using System E. Polar lipids extracted from Wildtype (WT), *pdtaS* knockout (Δms*pdtaS*), complemented (Δ:mt*pdtaS*) and phosphorylation defective (Δ:H303Q) strains and resolved on a 2D-TLC using solvent system E. Direction I; chloroform:methanol:water 60:30:60 v/v, direction II; chloroform:acetic acid:methanol:water 40:25:3:6 v/v. PIM – phosphatidylinositol mannoside (integers denote number of mannoside or acyl groups), PI – phosphatidyl inositol, PE – phosphatidyl ethanolamine, P – phospholipid. Representative image of duplicates (n = 2). **f**. Analysis of apolar FAMEs and MAMEs. Fatty acid methyl esters (FAMEs) and mycolic acid methyl esters (MAMEs) extracted from total apolar fraction separated on TLC plates using petroleum ether: ethyl acetate 95:5 v/v. **g**. Relative peak areas of lipid species detected in FAMEs and MAMEs preparation. Fold change in relative peak areas of lipid species in extracellular and intracellular apolar fractions normalized to wildtype peak areas have been plotted. C_n_ represents a fatty acid with a carbon chain of length n. 10Me – methyl group at C10. Dotted line represents the fold change of wildtype lipid species, set to 1. Quantitative analysis of TLC spot intensities for both polar and apolar lipid extracts can be found in Figure S4. GC-MS analyses of FAME extracts can be found in Figure S5 and Figure S6A. Physiological consequences of lipid loss can be found in Figure S6B and S6C.

The lipid loss identified by metabolomics was further characterized by analysis of the total lipids from cells grown in poor medium containing glycerol as the sole carbon source and supplemented with ^14^C-acetate to label all the lipids. Lipid extracts were then resolved on aluminium backed silica gel thin layer chromatography (TLC) plates and the bands were visualized by autoradiography. The total lipid extraction procedure allowed for the fractionation of lipids into two categories – apolar lipids and polar lipids.

Analysis of the polar lipid fraction was performed by 2D-TLC using System E (14), which allows separation and identification of the phosphatidyl ethanolamine (PE), phosphatidyl inositol (PI), phospholipid (P) and phosphatidylinositol mannosides (PIMs). Identification of individual lipids was performed based on comparison with previously published TLC analyses (15). Absence of functional PdtaS resulted in a reduction of polar membrane lipids such as PE, PI and P. In addition, early stage PIMs with two mannoside groups were unaffected but late stage hexamannoside PIM levels were also reduced (Fig. 3E and Fig. S4A).

Short chain fatty acids are usually produced by the fatty acid synthase I complex (FASI), which initiates the elongation of acetyl-CoA and malonyl-CoA units for bimodal production of C_18_ and C_26_ fatty acids in *M. smegmatis*. The products of FASI are modified and incorporated into the plasma membrane as phospholipids or are transferred to the fatty acid synthase II complex (FASII) for mycolic acid biosynthesis (16). In order to see if FASII products were similarly affected in the *pdtaS* knockout, lipid fractions were subjected to alkaline hydrolysis to produce fatty acid methyl esters (FAMEs) and mycolic acid methyl esters (MAMEs) which were then resolved using 1D-TLCs. While no changes were found in levels of MAMEs, FAMEs were reduced in Δms*pdtaS* and Δ:H303Q cells (Fig. 3F and Fig. S4B). The apolar lipid fraction was further fractionated into an apolar extracellular pool, consisting of non-covalently bound lipids on the outer surface and apolar intracellular pool, consisting of lipids extracted from cells after removal of the extracellular pool. FAMEs and MAMEs extracts of the apolar extracellular and intracellular fractions as well as the polar fraction were analyzed using GC-MS to identify and quantify the abundance of various FASI products. A marked reduction in the levels of C_18_, C_18:1_, C_24_ and C_26_ fatty acids in polar and apolar fractions was observed in cells lacking phosphorylation competent PdtaS. (Fig. 3G and Fig. S5-S6A). The presence of mycolates, which are made through elongation of shorter chain fatty acids from the FASI system indicates that both FASI and FASII lipid biosynthesis is taking place (since dysfunction of FASI will also disrupt FASII mediated mycolate formation). The loss of short chain fatty acids could therefore be attributed to increased lipid degradation in the knockout strains and not a dysfunction of lipid biosynthesis. The loss of membrane lipids also led to increased permeability to ethidium bromide (Fig. S6B) and increased susceptibility to streptomycin (Fig. S6C).

### Amino acid supplementation rescues growth and lipid loss

Since dipeptides are most commonly found as end products of protein degradation, we speculated that *pdtaS* knockout cells face amino acid starvation which results in a growth defect and breakdown of cellular lipids for salvage. Addition of cas amino acids, a common amino acid supplement should therefore rescue the growth defects and restore lipid levels to those of WT cells. As hypothesized, addition of cas amino acids rescued the growth defect of Δms*pdtaS* (Fig. 4A). Levels of FASI products were also restored in the *pdtaS* knockout upon cas amino acid supplementation (Fig. 4B and Fig. S7). These observations allowed us to propose a model of PdtaS-PdtaR function in the physiology of *M. smegmatis* (Fig. 4D). Briefly, we propose that nutrient depletion results in the production of an unknown ligand that activates PdtaS, resulting in signal transduction from PdtaS to PdtaR through phosphorylation. Phosphorylated PdtaR binds to target RNA to bring about changes in gene expression. In the case of Δms*pdtaS*, the absence of SK signalling should result in increased production and accumulation of the ligand due to the absence of ligand induced adaptation. The absence of PdtaS signalling may also induce the accumulation of the ligand due to the absence of a feedback loop, which is a common mechanism of regulation found in cellular metabolism.

**Figure 4.**
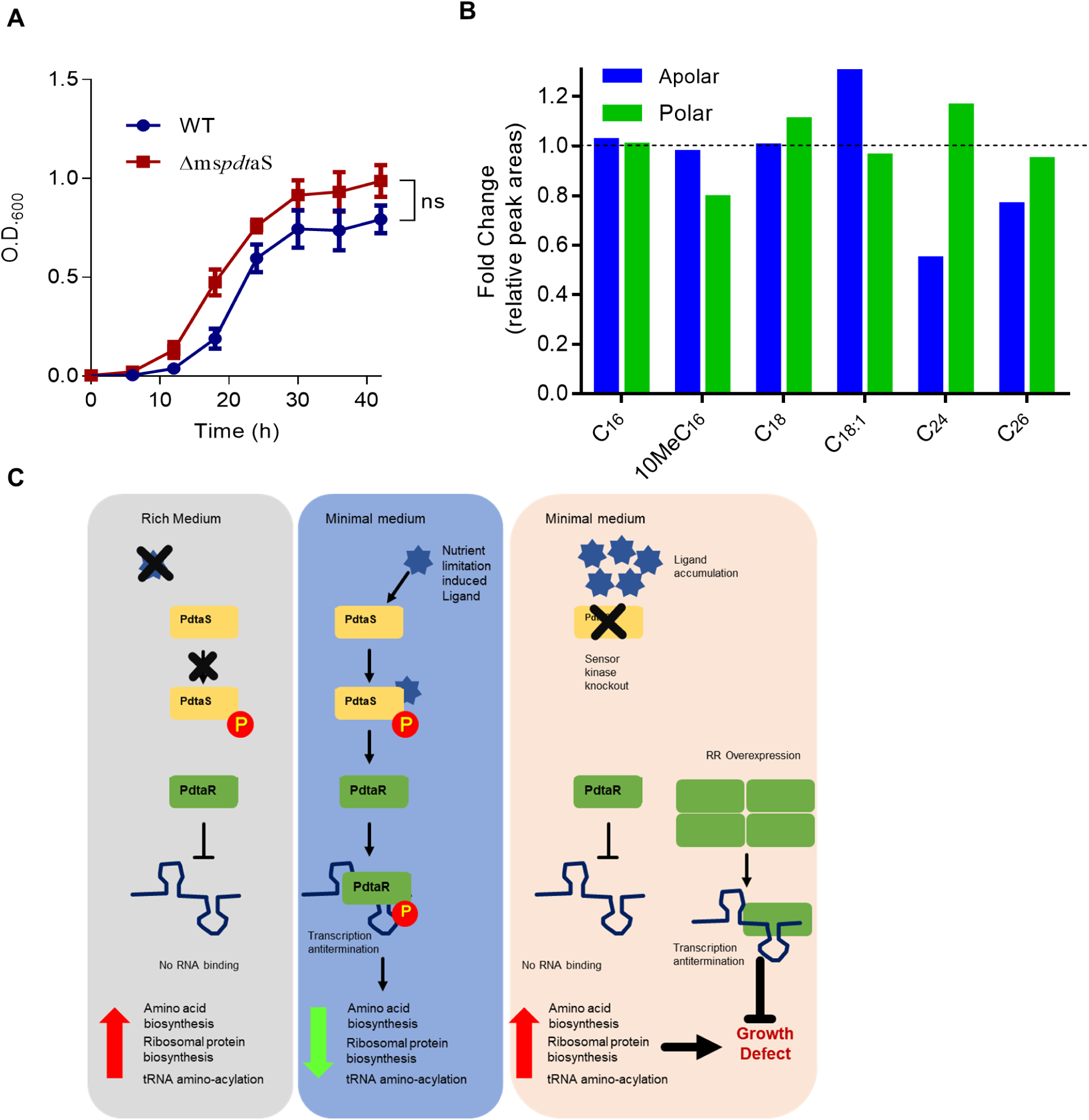
Amino acid supplementation rescues growth and lipid defects. a. Growth analysis of various M. smegmatis strains in presence of cas amino acid supplementation. Wildtype (blue) and Δms*pdtaS* (red) cells were inoculated in poor medium supplemented with 0.5% cas amino acids and the optical density of the cultures was monitored over time (n = 3). Error bars represent the standard error of the mean across replicates. **b**. Fold change in relative peak areas across all lipid fractions with cas-amino acids detected in total apolar (blue), and polar (green) fractions normalized to wildtype peak areas. 10Me – methyl group at C10. Dotted line represents the fold change of wildtype lipid species, set to 1 (n = 1). **c**. Model of PdtaS-PdtaR function in the physiology of *M. smegmatis*. The proposed model of PdtaS-PdtaR signalling involves the activation of PdtaS by a ligand during nutrient deprivation, resulting in activation of the signalling cascade and PdtaR mediated transcription anti-termination of target genes. If PdtaS is absent or inactive during nutrient deprivation, PdtaR mediated transcription anti-termination does not take place resulting in a growth defect. In such a case, it is hypothesized that the absence of PdtaS will result in the accumulation of the activating ligand. Furthermore, inducing target transcription anti-termination by overexpression of PdtaR in *pdtaS* knockout cells should relieve the growth defect. GC-MS analysis of FAME extracts after cas supplementation can be found in Figure S7.

### Cyclic diguanosine monophosphate binds to and activates PdtaS

With the above hypothesis, the comparative metabolome profiles of WT and Δms*pdtaS* lysates were scanned manually to find small molecules that accumualte in the knockout. One prospective small molecule, diguanosine tetraphosphate (pppGpG), was present at higher levels in Δms*pdtaS* (Fig. 5A). This molecule is an intermediate in the synthesis of the second messenger cyclic diguanosine monophosphate (c-di-GMP) from guanosine triphosphate (GTP) by the enzyme diguanylate cyclase (17). The accumulation of pppGpG made c-di-GMP a good candidate for the PdtaS GAF domain, which is well known for its ability to bind cyclic nucleotides (18). In addition, in silico PocketMatch screening of 28,533 non-redundant ligand containing binding pockets from PDB identified several potential hits comprising 61 unique ligands whose binding pockets matched that of PdtaS (Supplementary table 1). Among the top hits from PocketMatch was the structure of c-di-GMP bound LapD from *Pseudomonas fluorescence* (PDB ID: 3PJT) which further strengthened the probability that c-di-GMP could bind to PdtaS. C-di-GMP in Mycobacteria is known to regulate lipid transport and metabolism (19), multi-drug resistance (20), biofilm formation (21), cell wall polar lipid content (20) as well as dormancy and survival under nutrient deprivation (22). Many of these processes are perturbed in Δms*pdtaS* cells, therefore we tested for c-di-GMP binding to PdtaS using EGFP tagged PdtaS in a MicroScale Thermophoresis (MST) experiment. PdtaS-EGFP was confirmed to be enzymatically active and capable of autophosphorylation (Fig. S8A) and showed robust interaction with c-di-GMP with a K_D_ of 326 nM (Fig. 5B).

**Figure 5.**
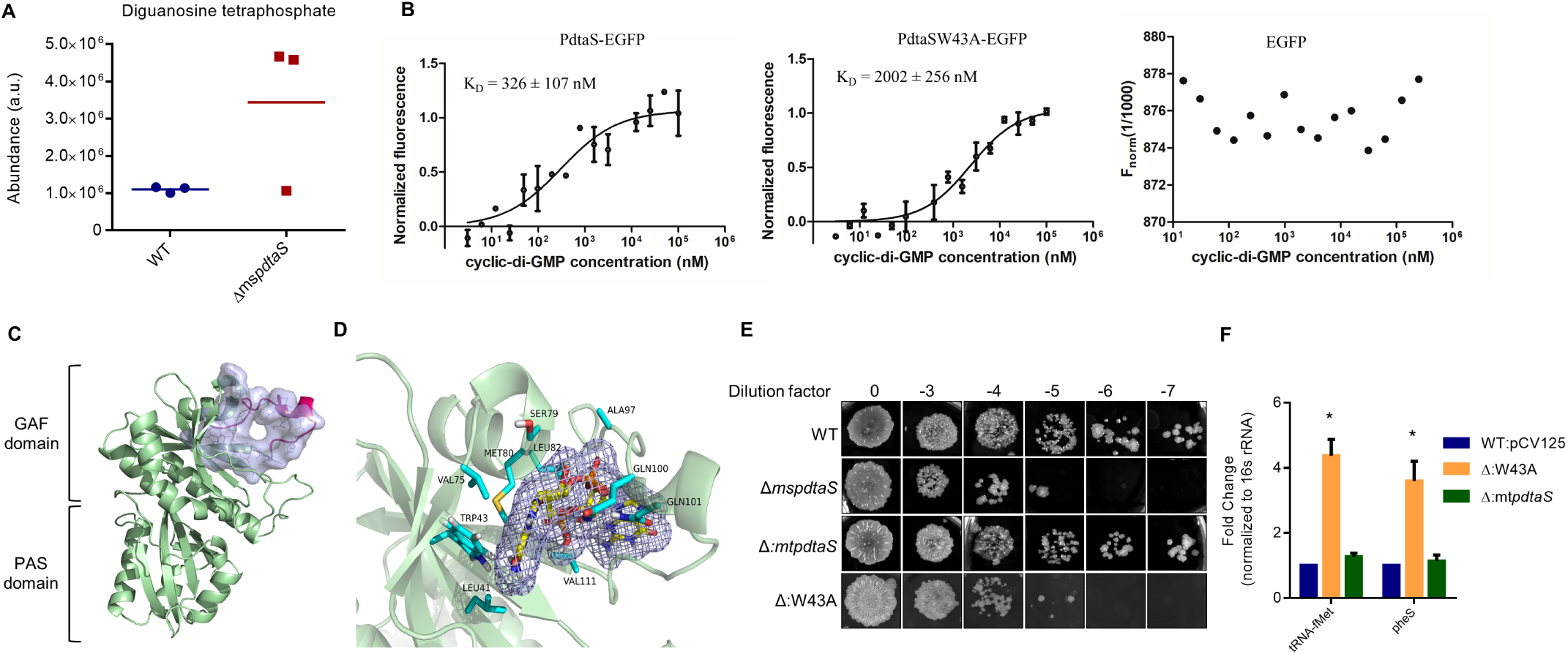
Cyclic diguanosine monophosphate binds to the PdtaS GAF domain. **a.** Diguanosine tetraphosphate accumulation in Δ*ms*pdtaS cells. Scatter plot of levels of diguanosine tetraphosphate in various strains (n = 3). **b**. Binding of c-di-GMP to PdtaS-EGFP, PdtaSW43A-EGFP and EGFP using MST. MicroScale thermophoresis of PdtaS-EGFP in the presence of increasing concentrations of cyclic-di-GMP. K_D_ = 326 +/- 107 nM for PdtaS-EGFP and 2002 +/- 256nM for PdtaSW43A-EGFP (n = 3). Validation of PdtaS-EGFP activity can be found in Figure S8A. **c.** Modelled protein structure of PdtaS using 3dba and 2ykf as templates. Modelled stretch of 18aa is shown in red while the consensus binding pocket in the GAF domain is represented in blue surface view with binding site residue in line representation. **d.** Docked c-di-GMP to the GAF domain pocket. Ligand is represented as yellow sticks with surface mesh while the interacting residues of the PdtaS GAF domain are in cyan sticks. Detailed interaction map of PdtaS ligand binding pocket and c-di-GMP can be found in Figure S8B. **e**. Complementation by mtPdtaS requires c-di-GMP binding. Dilutions of 36h cultures of wildtype (WT), Δ*pdtaS* and complemented strains Δ:mtp*dtaS* and Δ: W43A spotted onto agar plates. Representative of n = 3. **f.** Expression analysis of genes involved in translational processes in various strains as indicated. qRT-PCR expression analysis of tRNA-fMet and *pheS* in RNA isolated from wildtype *M. smegmatis*, Δ*pdtaS*, Δ:mt*pdtaS* and Δ: W43A strains. Expression is normalized using the 16s rRNA gene as an internal control. The fold change in gene expression for all strains relative to wildtype is plotted as mean of three independent experiments. Error bars represent standard error. The expression of genes in the wildtype is set to 1. * – p<0.05.

Since the crystal structure of the N-terminal domains of PdtaS (2ykf) was available, we decided to see if the GAF domain can bind to c-di-GMP (C2E) through molecular docking. However, the accurate docking of c-di-GMP onto the crystal structure of PdtaS required refining of the GAF domain lid, which was unstructured and hypothesized to attain a stable secondary structure upon ligand binding (23). We therefore modelled the unstructured portion using 3dba_A, a template identified through PDBeFold, to predict a likely conformation for the unstructured region (Fig. 5C). Docking revealed that c-di-GMP was capable of strong binding to monomeric PdtaS, with one of the two rings of c-di-GMP stabilized by a tryptophan residue (Trp^43^) similar to the stabilization of c-di-GMP by CckA of *Caulobacter crescentus* (24) (Fig. 5D). The docking of c-di-GMP to the GAF ligand binding pocket also provided details of stabilizing interactions between the GAF pocket residues and c-di-GMP (Fig. S8B). In order to see if c-di-GMP binding is essential for PdtaS mediated adaptation to nutrient deprivation, the Trp^43^ residue was mutated to Ala in the complementation plasmid as well as in the N-terminal EGFP tagged construct for protein purification. Analysis of the c-di-GMP binding mutant using MST revealed a ten-fold decrease in binding affinities compared to the wildtype protein (Fig. 5B). The complementation construct carrying the W43A mutation was electroporated into *Δ*ms*pdtaS* to create Δ:W43A strain.

The mutant Δ:W43A PdtaS was unable to rescue the growth or transcriptional dysregulation of *pdtaS* knockout cells (Fig. 5E and 5F), indicating that c-di-GMP is indeed the ligand for PdtaS binding to c-di-GMP is essential for its function during conditions of nutrient deprivation. It is interesting that a ten-fold decrease in binding affinities results in a complete loss of complementation, indicating the narrow range of c-di-GMP concentrations at which PdtaS functions.

### A novel interactome underlies PdtaS-PdtaR mediated adaptation

Transcriptomic and metabolomic analysis of the *pdtaS* knockout strain identified large scale changes in amino acid metabolism and translation, indicated by the strong induction of amino acid biosynthesis, tRNA synthetases and 50s as well as 30s ribosomal protein genes (Fig. S9A). The cellular requirement for amino acids was underscored by the accumulation of proteolytic dipeptides and the recovery of growth with the addition of cas amino acids. In addition, both polar and apolar lipid levels were depleted in PdtaS deficient cells. Again, the lipid loss was recovered upon addition of cas amino acids to the medium. This indicated a link between the translation machinery, amino acid biosynthesis and lipid homeostasis. However, to identify key processes underlying and connecting perturbations in these diverse metabolic pathways, a systems level understanding was needed. Towards this, we constructed a knowledge-based *M*. *smegmatis* protein-protein-interaction network (PPI-net) that represents a global connectivity map of *M. smegmatis* as described in the methods section. The network consists of 80,836 interactions (edges) among 5505 proteins (nodes), thus covering 82% of *M. smegmatis* proteome. The network was made condition specific by integrating gene expression data in the form of node and edge weights and then mined using a sensitive network mining algorithm as described earlier (25) to identify the top-response network perturbed in the *pdtaS* knockout. These paths represent the processes that are taking place differently in the mutant strain as compared to the WT and is referred to as the top-response network. The top-response network, thus generated consists of 440 nodes with 473 interactions out of which 192 belong to intermediary metabolism and respiration and 104 to lipid metabolism (Fig. 6A). This reinforces previous work which identified that a *M. smegmatis pdtaS* knockout is susceptible to aminoglycosides and respiratory chain inhibitors (8). From the top response network, the arginine biosynthetic operon was found to transmit perturbations to the holo-(acyl-carrier-protein) synthase (AcpS – MSMEG_4756), the ribosomal protein operon (RpsL) and tRNA synthetases PheS and PheT. PheS and PheT were found to link to fatty acid synthase I (Fas I – MSMEG_4757). These major hubs were involved in further transmitting perturbations to downstream nodes involved in lipid metabolism, intermediary metabolism, respiration, information pathways as well as cell wall and cell wall processes. ClueGO enrichment of the top response network revealed intermediary metabolism and respiration genes as the largest perturbed processes, followed by lipid metabolism, information pathways and cell wall and cell processes (Fig. S9B).

**Figure 6.**
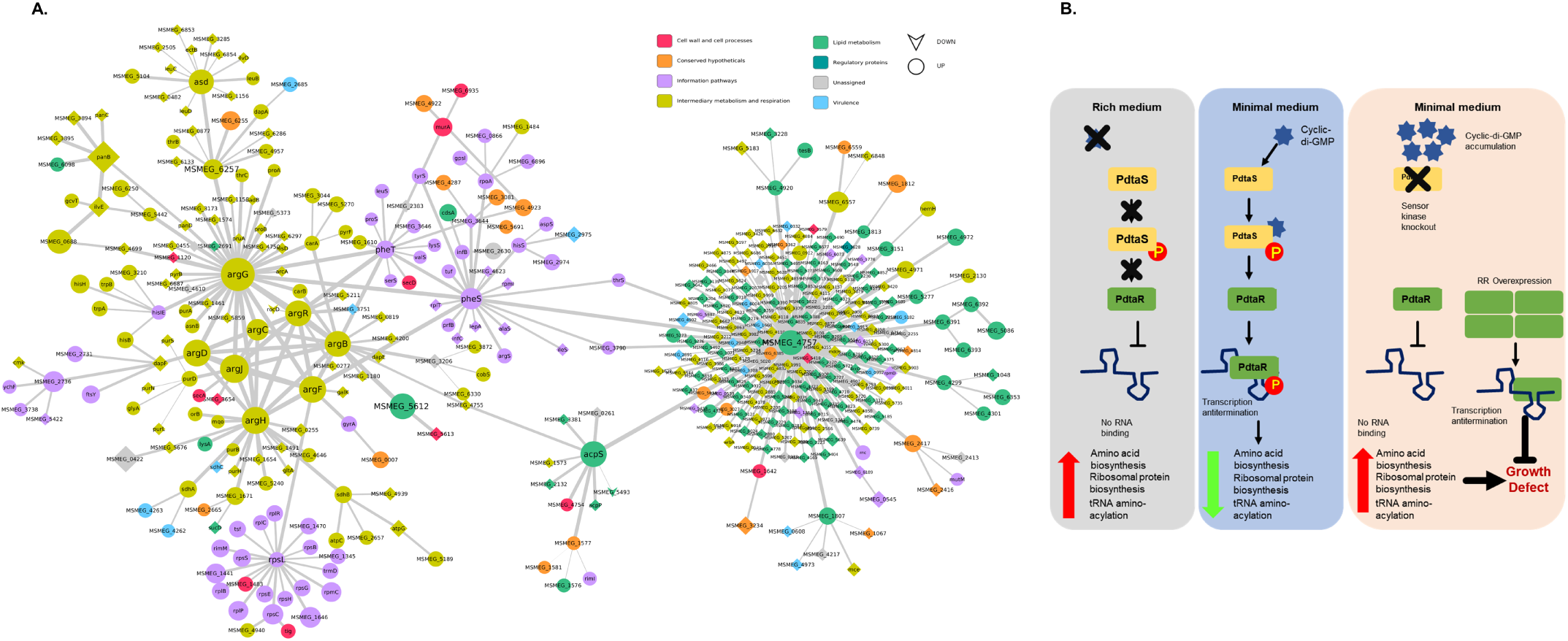
Protein-Protein interaction map of PdtaS deficient cells and Schematic Representation of PdtaS-PdtaR signalling pathway. **a.** Top-response network of *pdtaS* knockout strain compared to wildtype. Proteins are represented as nodes and interactions between them as edges. Nodes are coloured based upon Mycobrowser functional categories mapped from Mtb homologs and the size of each nodes is based upon fold change value of gene expression. Significantly up and down regulated genes are represented as circles and inverted triangles respectively while the non-differentially expressed genes are represented as diamonds. Heatmap of genes from the microarray data that are representative of the protein-protein interaction network can be found in Figure S9A. Frequency bar plot of gene functional categories represented in the protein-protein interaction network can be found in Figure S9B. **b.** During conditions of nutritional abundance, cellular biosynthetic and translational processes are upregulated to facilitate utilization of nutrients and rapid growth, with PdtaS-PdtaR signalling absent (left Panel). Nutrient paucity (centre panel) results in increasing intracellular concentrations of c-di-GMP which binds to the sensor kinase PdtaS and promotes PdtaS-PdtaR signalling resulting in transcriptional adaptation to nutrient deprivation. Removal of PdtaS or prevention of PdtaS signalling during nutrient deprivation (right panel) results in transcriptional dysregulation of biosynthetic and translational processes, resulting in a growth defect and absence of mycobacterial adaptation.

## Discussion

Two component systems of *Mycobacterium tuberculosis* have recently emerged as promising drug targets in anti-tuberculosis chemotherapy (2, 3, 26, 27), with several TCS RRs implicated in key adaptive responses such as response to pH, envelope stress, phosphate limitation and hypoxia (5). However, the mechanistic details of TCS signalling, especially the role of SKs and their ligand binding domains in modulating responses to these conditions are yet to be elucidated. Indeed, the absence of knowledge about SK ligands is a limitation common to TCSs across bacterial species. It is often known that a particular TCS is important for adaptation to a broad condition such as oxidative stress or nutrient limitation, but ligands that bind to and activate an SK remain elusive (28). Additionally, due to the great diversity of signals sensed by TCSs despite having a fixed set of domain architectures in their sensory domains, the nature of ligands that bind to the SKs are not readily apparent from the structure of their sensory domains (29). Currently, only two SKs DevS and SenX3 have known ligands and characterized mechanisms of activation (30, 31). This is essential as in order to identify the ideal TCS drug target, the details about what TCS SKs sense and how signal transduction occurs need to be elucidated.

Here we report the first comprehensive characterization of a relatively unknown TCS, the PdtaS-PdtaR system – from stimulus to response (Fig. 6B). It was first reported as a bona fide TCS in 2005 (10), the role of PdtaS in antibiotic resistance was only reported recently (8) and crystal structures for both SK and RR (12, 23) are also available. Here we characterize the physiological function of PdtaS-PdtaR signalling using transcriptomic and metabolomic analysis and demonstrate that this TCS plays an essential role in Mycobacterial adaptation to poor nutrient conditions. Absence of PdtaS in *Mycobacterium smegmatis* leads to global transcriptional dysregulation of biosynthesis and transmembrane transport, leading to a loss of lipids and increased membrane permeability. We show that targeting this TCS by prevention of PdtaS phosphorylation or PdtaR RNA binding will result in poor growth, loss of polar and apolar lipid content and amino acid starvation. Additionally, we have identified c-di-GMP as a ligand and activator of PdtaS autophosphorylation and have characterized the binding pocket and interacting residues. The promoter of the *M. smegmatis* diguanylate cyclase, involved in c-di-GMP synthesis and degradation, is induced during starvation and repressed upon addition of glucose (32). Moreover, the diguanylate cyclase is essential for long-term survival under nutrient starvation (33) indicating that the PdtaS-PdtaR TCS is important for regulating genes that help Mycobacteria adapt to low nutrient conditions. Our in silico network analysis also revealed the arginine biosynthesis operon as a centrally perturbed node during PdtaS deficiency. Members of this operon have been characterized as essential for Mtb survival and virulence (34) and have been proposed as targets for anti-tuberculosis chemotherapy (35, 36).

Our characterization, supplemented with existing observations that PdtaR is constitutively expressed during macrophage infection (7), highly expressed during infection in SCID mice (37) and is a key secreted antigen and potential vaccine candidate (38) strengthens the case for the exploration of PdtaS-PdtaR inhibitors as a drug target. Development of c-di-GMP mimetics as PdtaS inhibitors also promise to affect a wider range of Mycobacterial functions such as lipid transport and metabolism (19), multi-drug resistance, biofilm formation, cell wall polar lipid and glycopeptidolipid content (20, 21, 39) and dormancy and survival under nutrient deprivation (22, 33), reducing the chances of drug resistance through target mutation. Overall, we have unravelled the details of PdtaS-PdtaR signalling, added to our knowledge of ligands for Mycobacterial SKs and uncovered a promising target for future anti-tubercular therapy.

## Materials and Methods

### Materials

All bacterial growth media, growth supplements, fine chemicals, amino acids and cas amino acids were purchased from Sigma-Aldrich (U.S.A.). Antibiotics and dithiothreitol (DTT) were purchased from Toku-E (Japan). Restriction endonucleases, T4 DNA ligase, DNAseI, RNAse A, DNA and protein markers were purchased from Thermo Fisher Scientific (U.S.A.). Ni Sepharose™ 6 Fast Flow resin was purchased from GE Healthcare (U.S.A.). All primers were synthesized and sequencing performed by Bioserve (India). γ^32^P ATP (>3500 Ci/mmol) was purchased from BRIT-Jonaki (India).

### Bacterial strains and growth conditions

All cloning steps were carried out in *Escherichia coli* DH10β (Thermo Fisher Scientific). For protein purification, the expression strain ArcticExpress (DE3) (Agilent Technologies) was used. All *E.coli* were grown in Luria-Bertani (LB) medium with appropriate antibiotics at 37°C with shaking at 150rpm. Antibiotics were used at the following concentrations – ampicillin 100mg/l, kanamycin 25mg/l, hygromycin 100mg/l.

*Mycobacterium smegmatis* mc^2^155 was grown in Middlebrook 7H9 Broth Base (HiMedia) or Tryptone Soya Broth (HiMedia) at 37°C with shaking at 150rpm. 7H9 was supplemented with 0.025% tyloxapol and 0.2% glycerol unless otherwise specified.

### Gene cloning and site directed mutagenesis

The gene coding for PdtaS from *M. smegmatis* mc^2^155 and *M. tuberculosis* H37Rv were PCR amplified using isolated genomic DNA as a template, Pfu polymerase and gene specific primers containing appropriate restriction sites at their ends. PCR amplified genes were cloned into appropriate vectors for *E. coli* and mycobacterial expression. All constructs were verified by sequencing andSsite directed mutagenesis for the generation of point mutations as well as domain deletions were performed using Quick Change Site-Directed Mutagenesis protocol using Phusion polymerase (New England BioLabs, USA). Mutations/deletions introduced were verified by DNA sequencing.

### RNA isolation and quantitative reverse transcription PCR

RNA was isolated from mid-log phase cultures of *M. smegmatis* using the RNeasy Mini Kit (Qiagen, USA) according to the manufacturer’s recommended protocol. Briefly, 10 ml of mid-log phase cells were harvested and resuspended in 1ml of QIAzol Lysis Reagent followed by cell lysis using the Qiagen TissueLyser II. Lysis was performed by bead beating using 0.1mm Zirconia/Silica beads (Thomas Scientific, USA) in 5 cycles 45s each with 30s intervals. After bead beating, RNA was precipitated with ethanol and loaded onto the purification columns. Columns were washed with ethanol containing wash buffers followed by elution of the RNA. Eluted RNA was treated with DNAseI (Thermo Fisher Scientific, USA) to remove any DNA contamination, followed by column clean-up according to the Qiagen clean-up protocol.

Quantification of RNA yield was performed using the NanoDrop® ND-1000 UV-Vis Spectrophotometer (NanoDrop Technologies, USA). 1µg of purified RNA was used to make cDNA using the iScript™ cDNA synthesis kit (Bio-Rad, USA) using the manufacturer’s recommended protocol. 1µl of the cDNA reaction was used to set up qRT-PCR using the iTaq™ Universal SYBR® Green Supermix in the Rotor-Gene Q (Qiagen, USA) according to the manufacturer’s protocol.

### Autophosphorylation and phosphotransfer assays

Purified SKs (150pmol unless stated otherwise) were incubated in kinase buffer (50mM tris-Cl pH 8, 50mM KCl, 10mM MgCl_2_) containing 1µCi of [γ^32^P] ATP for 60min or for time points specified at 30°C. Reactions were terminated by the addition of 1X SDS-PAGE loading buffer (2% SDS, 50mM tris-Cl pH 8, 0.02% bromophenol blue, 1% β-mercaptoethanol, 10% glycerol) and loaded onto the wells of an SDS-PAGE gel. Samples were resolved on a 15% SDS-PAGE gel, washed with distilled water and exposed to a phosphor screen (Fujifilm Bas cassette) overnight followed by imaging with a Typhoon 9210 phosphorimager (GE Healthcare).

For the effect of c-di-GMP on autophosphorylation, 150pmol of PdtaS was mixed with various concentrations of c-di-GMP and incubated in kinase buffer containing [γ^32^P] ATP. In order to study the phosphotransfer from SK to RR, SK∼P was prepared as described above, following which 150pmoles of purified RR diluted in 1X kinase buffer was added to the reaction. The reaction was incubated at 30°C for 15min unless otherwise specified followed by reaction termination by the addition of 1X SDS-PAGE loading buffer. The proteins were resolved on 15% SDS-PAGE gels followed by autoradiography.

Densitometric analysis of autoradiograms was performed using ImageJ.

### Growth curves and dilution spot assays

Primary cultures of *M. smegmatis* strains were grown in TSB (unless otherwise specified) for 24h or until the O.D. reached 0.8-1.0, after which they were sub-cultured at a starting O.D. of 0.005 in the medium in which the experiment was carried out. For starvation experiments, primary cultures were washed thrice with 7H9 alone (no carbon source, no supplements) followed by resuspension in 7H9 alone for 8h, following which they were sub-cultured at a starting O.D. of 0.005 in the medium used for the growth curve.

At specified time-points, 200µl of cultures were pipetted into the wells of flat, transparent clear bottom 96-well plates (Corning, USA) in triplicate and the absorbance of the wells were measured at 600nm using the bottom reading mode of the Infinite M1000 PRO multimode fluorescence plate reader (Tecan). The absorbance measurements were plotted as a function of time to obtain a growth curve for the culture.

A single time point assay for the growth of *M. smegmatis* was standardized using spot dilutions on tryptone soy agar (TSA) (HiMedia) plates. Briefly, primary cultures of *M. smegmatis* strains were inoculated in tubes containing the appropriate medium at an O.D. of 0.005. Cultures were allowed to grow for 30h (mid-log phase for wildtype cells) following which serial ten-fold dilutions of cultures were made in 7H9 base medium. 20µl of each dilution was spotted carefully on the surface of TSA plates containing appropriate antibiotics. Spots were allowed to dry (no visible moisture on the agar surface) following which they were incubated at 37°C for 2-3 days to allow for growth of single colonies.

Plates were imaged using a conventional camera and dilution spot pictures were collated as an image to enable comparison between strains.

### MicroScale Thermophoresis

MicroScale Thermophoresis (MST) experiments were performed using the Monolith NT.115 system (NanoTemper Technologies, Germany). The laser power was set at 20% and the LED at 100%. Equilibrated mixtures containing 5µM of PdtaS-EGFP and varying concentrations of c-di-GMP were loading onto hydrophobic capillaries (NanoTemper Technologies). The laser on and off times were set at 30s and 5s respectively. Normalized fluorescence change was plotted against concentration of c-di-GMP to get the binding curve. Curve fitting using NT analysis software was used to find K_D_ values.

#### Statistical Analyses

Statistical analyses for significance were performed using Student’s t-test unless otherwise mentioned. For all experiments, number of independent biological replicates used are indicated by ‘n’.

## Acknowledgements

We would like to thank Dr. Dipankar Chatterjee, Dr. Umesh Varshney and Dr. Vandana Malhotra for their inputs and expertise. We also thank Dr. Eliza Peterson for their inputs and critical reading of this manuscript. Dr. Warwick Dunn mass spectrometry facility, University of Birmingham is acknowledged for MS/MS analysis.

## Author contributions

VNH, designed the study, performed the experiments, analyzed the data and wrote the manuscript; CT, performed the docking studies; AS, generated the knockout strains; RG, performed the site directed mutagenesis; DPS and GDS, performed the MST experiments; NC and AB, designed the study, analyzed the data and DKS, conceived the study, analyzed the data and wrote the manuscript.

## Supplementary Methods

### Whole genome microarray and analysis

#### RNA quality and quantity assessment

The isolated RNA was quantified using a NanoDrop spectrophotometer (ND-2000, Thermo fisher scientific, USA). The quantity was measured by absorbance at 260nm and the purity was assessed using 260/230 and 260/280 for salt/organic solvents and protein/phenol/other contaminants respectively. The isolated total RNA integrity was analyzed with an Agilent 2100 Bioanalyzer (Agilent Technologies, Palo Alto, CA) according to the manufacturer’s instructions. The 18S and 28S rRNA ratio was obtained from 2100 Expert software (Agilent Technologies) and RNA integrity number was obtained from RIN Beta Version Software (Agilent Technologies) was calculated.

#### Labelling and microarray hybridization

The microarray hybridization and scanning were performed at the Agilent certified microarray facility of Genotypic Technology, Bengaluru, India. The samples for Gene expression were labeled using Agilent Quick-Amp labeling Kit (p/n5190-0442). In brief, 500ng of total RNA were reverse transcribed at 40°C using random primer tagged to a T7 polymerase promoter and converted to double stranded cDNA. Synthesized double stranded cDNA were used as template for cRNA generation. cRNA was generated by in vitro transcription and the dye Cy3 CTP (Agilent) was incorporated during this step. The cDNA synthesis and in vitro transcription steps were carried out at 40°C. Labeled cRNA was cleaned up using Qiagen RNeasy columns (Qiagen, Cat No: 74106) and quality assessed for yields and specific activity using the Nanodrop ND-2000. 600ng of labeled cRNA sample were fragmented at 60°C and hybridized on to an Agilent *M. smegmatis* Gene Expression Microarray 8X15K. Fragmentation of labeled cRNA and hybridization were done using the Gene Expression Hybridization kit (Agilent Technologies, In situ Hybridization kit, Part Number 5190-6420). Hybridization was carried out in Agilent’s Surehyb Chambers at 65° C for 16 hours. The hybridized slides were washed using Agilent Gene Expression wash buffers (Agilent Technologies, Part Number 5188-5327) and scanned using the Agilent Microarray Scanner (Agilent Technologies, Part Number G2600D).

#### Microarray data analysis

Microarray data preprocessing and normalization was performed in R using limma, a part of the Bioconductor package (58). Background correction was performed using normexp() after the extraction of median signal and background intensities using read.maimages() function. Adjusted signals were then log_2_ transformed and quantile normalised to make the intensity distributions similar across samples. Differential gene expression analysis between mutant and WT condition was performed using ebayes() and genes with log_2_ FC ≥1≤ −1 and p-value < 0.05 were considered to be significantly diffrentially expressed (DEGs). A functional enrichment analysis for the DEGs was performed with ClueGOv2.3.5, cytoscape plugin (59).

We acknowledge Genotypic Technology Private Limited Bangalore for the microarray processing reported in this publication. Data and details of the gene expression microarray have been deposited in the Gene Expression Omnibus (GEO) repository under accession ID GSE108559.

#### UPLC-MS: Sample preparation

20ml of mid-log phase cultures of M. smegmatis was quenched with an equal volume of methanol:water 60:40 v/v solution that was pre-chilled at −60°C, followed by immediate centrifugation to harvest the cells. The pellets were then transferred to pre-weighed centrifuge tubes using 0.5-1ml of PBS, following by centrifugation to remove the PBS.

The wet-weight of the cell pellets were measured and 4µl/mg of methanol and 0.85µl/mg of water was added to resuspend the cells followed by bead beating as described above (section 2.6). The homogenized mixtures were transferred using a glass Pasteur pipette to glass tubes, followed by the addition of 1µl/mg of methanol and 0.9µl/mg water. The mixtures were vortexed for 15s to suspend any remaining material. The tubes containing the mixtures were then placed on ice followed by the addition of 5µl/mg chloroform and 2µl/mg water. The tubes were vortexed for a further 30s and then incubated on ice for 30s. The tubes were then centrifuged at 1800g for 10min at 4°C and then allowed to stand still at room temperature for 5min. The upper polar layer and the lower apolar layer were separated and transferred to fresh glass vials using glass Pasteur pipettes. The samples were lyophilized and later analysed using Ultra Performance Liquid Chromatography – Mass Spectrometry analysis.

For the analysis of water-soluble metabolites, 100 µL of acetonitrile (LC-MS grade, LiChrosolv, Merck) was added to dried sample followed by vortex mixing (15 seconds), centrifugation (13,000×g, 15 min) and transfer of the 90 µL clear supernatant to a glass LC autosampler vial (VI-04-12-02RVG 300μl Plastic, Chromatography Direct, UK). For the analysis of lipid metabolites, 100 µL of mixture isopropyl alcohol IPA (LC-MS grade, LiChrosolv, Merck) was added to the dried sample followed by vortex mixing (15 seconds), centrifugation (13,000×g, 15 minutes) and transfer of the 90 µL clear supernatant to a glass LC autosampler vial (VI-04-12-02RVG 300μl Plastic, Chromatography Direct, UK).

#### UPLC-MS: Data acquisition

The samples were analysed applying two Ultra Performance Liquid Chromatography-Mass Spectrometry (UPLC-MS) methods using a Dionex UltiMate 3000 Rapid Separation LC system (Thermo Fisher Scientific, MA, USA) coupled with and electrospray Q Exactive Focus mass spectrometer (Thermo Fisher Scientific, MA, USA). Polar extracts were analysed on a Accucore-150-Amide-HILIC column (100 × 2.1 mm, 2.6 μm, Thermo Fisher Scientific, MA, USA). Mobile phase A consisted of 10 mM ammonium formate and 0.1% formic acid in 95% acetonitrile/water and mobile phase B consisted of 10 mM ammonium formate and 0.1% formic acid in 50% acetonitrile/water. Flow rate was set for 0.50 mL·min-1 with the following gradient: t=0.0, 1% B; t=1.0, 1% B; t=3.0, 15% B; t=6.0, 50% B; t=9.0, 95% B; t=10.0, 95% B; t=10.5, 1% B; t=14.0, 1% B, all changes were linear with curve = 5. The column temperature was set to 35 °C and the injection volume was 2 μL. Data were acquired in positive and negative ionisation modes separately within the mass range of 70 – 1050 m/z at resolution 70,000 (FWHM at m/z 200). Ion source parameters were set as follows: Sheath gas = 53 arbitrary units, Aux gas = 14 arbitrary units, Sweep gas = 3 arbitrary units, Spray Voltage = 3.5kV, Capillary temp. = 269 °C, Aux gas heater temp. = 438 °C. Data dependent MS2 in ‘Discovery mode’ was used for the MS/MS spectra acquisition using following settings: resolution = 17,500 (FWHM at m/z 200); Isolation width = 3.0 m/z; stepped normalised collision energies (stepped NCE) = 25, 60, 100%. Spectra were acquired in three different mass ranges: 50 – 200 m/z; 200 – 400 m/z; 400 – 1000 m/z.

Non-polar extracts were analysed on Hypersil GOLD column (100 × 2.1mm, 1.9 µm; Thermo Fisher Scientific, MA, USA). Mobile phase A consisted of 10 mM ammonium formate and 0.1% formic acid in 60% acetonitrile/water and mobile phase B consisted of 10 mM ammonium formate and 0.1% formic acid in 90% propan-2-ol/water. Flow rate was set for 0.40 mL.min-1 with the following gradient: t=0.0, 20% B; t=0.5, 20% B, t=8.5, 100% B; t=9.5, 100% B; t=11.5, 20% B; t=14.0, 20% B, all changes were linear with curve = 5. The column temperature was set to 55 °C and the injection volume was 2μL. Data were acquired in positive and negative ionisation mode separately within the mass range of 150 – 2000 m/z at resolution 70,000 (FWHM at m/z 200). Ion source parameters were set as follows: Sheath gas = 50 arbitrary units, Aux gas = 13 arbitrary units, Sweep gas = 3 arbitrary units, Spray Voltage = 3.5kV, Capillary temp. = 263 °C, Aux gas heater temp. = 425 °C. A Thermo ExactiveTune 2.8 SP1 build 2806 was used as an instrument control software in both cases and data were acquired in profile mode.

#### UPLC-MS: Raw data processing

Raw data acquired in each analytical batch were converted from the instrument-specific format to the mzML file format applying the open access ProteoWizard software (60).

Deconvolution was performed for each assay separately with XCMS software according to the following settings of Min peak width (4 for HILIC and 6 for lipids); max peak width (30); ppm (12 for HILIC and 14 for lipids); mzdiff (0.001); bandwidth (0.25); mzwid (0.01) (61, 62). A data matrix of metabolite features (m/z-retention time pairs) vs. samples was constructed with peak areas provided where the metabolite feature was detected for each sample.

Extracted peak tables for each assay were used to assess quality of the data sets. In particular data were checked for possible retention time and m/z drifts, and signal intensity drifts related to the sample measurement order.

Putative annotation of metabolites or metabolite groups was performed by applying the PUTMEDID-LCMS workflows operating in the Taverna workflow environment (63). We applied 12 ppm mass error for HILIC and 14 ppm mass error for lipids with a retention time range of 2 s in feature grouping and molecular formula and metabolite matching. Because different metabolites can be detected with the same accurate m/z (for example, isomers with the same molecular formula), multiple annotations could be observed for a single detected metabolite feature. Also, a single metabolite could be detected as multiple molecules, particularly as a different type of ion (e.g., protonated and sodiated ions). Throughout this article, the term “metabolite” refers to either single metabolites or groups of molecules with the same retention time and the same accurate m/z.

#### UPLC-MS: Statistical analysis

All data were sample normalised to total peak area using the following calculation:

Normalised peak area (metabolitex, sampley) (%) =

(Peak Area (metabolitex) / Total peak area for all metabolites for sampley)) × 100

Mann-Whitney U tests were applied to identify statistically significant metabolite features (critical p-value<0.05). Fold changes were calculated using the median of responses for all samples in each of the two classes:

Fold change (median, class A) / (median, class B)

Finally, biologically important metabolites were classified according to metabolite class applying a manual process to investigate if specific metabolite classes were enriched.

#### Lipid extraction

All glass tubes for lipid extraction were pre-weighed. All centrifugations performed at room temperature, 3500rpm for 5min. All incubations and extractions were also performed at room temperature. 10ml of labelled cells were pelleted, transferred to a glass tube labelled **tube A** (16×100mm with a Teflon lined stopper in the lid) and dried under nitrogen until no visible moisture remained. To the dried pellet was added 2ml of methanol:0.3% NaCl (10:1) followed by 1ml of petroleum ether (60-80°C). The mixture was then placed on a rotator for 15min followed by centrifugation. The upper layer was transferred to a separate tube (**tube B)**. To tube A was added another 2ml of petroleum ether, mixed for 15min followed by centrifugation. The upper layer of this step was transferred to **tube B** which now contains the supernatant from two extractions of the cell pellet. **Tube B** contains apolar lipids and was dried under a stream of nitrogen and weighed. If extracellular and intracellular apolar lipids were to be isolated separately, two petroleum ether extractions (2ml each) prior to the addition of methanol:0.3% NaCl were performed as described above, and transferred to a separate tube, dried and weighed. The continuation of the above protocol after pre-extraction with petroleum ether will yield the intracellular apolar lipid fraction.

To **tube A** after apolar lipid extraction, 2.3ml of chloroform:methanol:0.3% NaCl (9:10:3) was added and mixed with the remaining pellet on a rotator for 60min followed by centrifugation. The supernatant was transferred to **tube C**. To **tube A** was added 750µl of chloroform:methanol:0.3% NaCl (5:10:4) and the mixture was incubated at on a rotator for 30min followed by centrifugation. The supernatant was transferred to **tube C** containing the supernatant from the first extraction. A second round of 5:10:4 extraction was performed and the supernatant pooled into **tube C**. To **tube C**, now containing the supernatants from three extraction steps was added 1.3ml of chloroform + 1.3ml of 0.3% NaCl. This resulted in the formation of two phases. **Tube C** was rotated for 5min followed by centrifugated.

The upper layer of **tube C** was discarded, careful not to allow mixing of the two layers or contamination of the lower layer with upper layer contents. The lower layer was transferred to **tube D**. **Tube D** contains polar lipid extracts and was dried under a stream of nitrogen and weighed.

In order to extract Fatty acid and mycolic acid methyl esters (FAMEs and MAMEs) from apolar and polar fractions extracted previously, 2ml of 5% tetrabutylammonium hydroxide was added to the cell pellet/dried lipids and incubated overnight at 100°C using a heat block. This step results in the alkaline hydrolysis of FAMEs and MAMEs. Following alkaline hydrolysis, the tubes were allowed to cool down to room temperature methyl esterified by the sequential addition of the following chemicals: 4ml of dichloromethane, 300µl of iodomethane and 2ml water. The tube was mixed for 1h on a rotator followed by centrifugation. The upper aqueous phase was discarded and 4ml of water was added to the lower organic phase (water wash) and incubated on a rotator for 15min followed by centrifugation and discard of the aqueous supernatant. The water wash was repeated two more times. The lower organic phase was dried under a stream of nitrogen until no visible moisture remained (usually takes at least 30min).

To the dried residue was added 3ml of diethylether followed by sonication in a water bath sonicator for 10min at room temperature. Sonication was repeated until a uniform suspension was obtained. The mixture was centrifuged and the upper layer containing FAMEs and MAMEs was transferred to a separate tube, dried and weighed.

#### Lipid analysis

Dried lipid extracts were resuspended in chloroform:methanol 2:1 v/v and used for scintillation counting to estimate radioactivity. Appropriate volumes corresponding to equal radioactive counts (10,000 cpm) were spotted onto aluminium backed silica TLC plates (5554 silica gel 60F524; Merck) and run in solvent system E (Direction I; chloroform:methanol:water 60:30:60 v/v, direction II; chloroform:acetic acid:methanol:water 40:25:3:6 v/v). FAMEs and MAMEs spotted on TLC plates were run using petroleum ether/acetone 95:5 v/v as the mobile phase.

Lipids were visualized by overnight exposure to Kodak X-Omat AR film.

2mg of dried FAMEs and MAMEs from unlabelled cells were submitted for GC-MS analysis of lipid peaks.

#### Construction of a response network

Global protein-protein interaction network of Msm was constructed using STRING v10 (64) database. A combined score cutoff of 700 was used to extract high confidence interactions, which captures both physical as well as functional associations among interacting proteins. The final constructed network had 5505 proteins with 80,836 interactions among them. In the network, proteins are represented as nodes and interactions among them as edges. This basal network was made context specific by integrating gene expression data onto the PPI network by assigning weights to nodes and edges (65) calculated as follows:

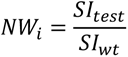

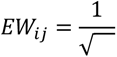

where, NW_i_ is the node weight of node *i*, SI_test_ and SI_wt_ is the signal intensity of gene *i* in test and wild type condition respectively and EW_ij_ is the edge weight between node i and j.

To identify perturbations in the given condition, (as compared to its control), we used an in-house sensitive network mining algorithm which (a) computes all-to-all shortest paths in the weighted network using Djikstra’s algorithm (implemented in the Zen library (http://www.networkdynamics.org/static/zen/html/api/algorithms/shortest_path.html), (b) computes path scores by taking a summation of all edge scores in that path and (c) normalizes and ranks them to identify the most perturbed paths and constitutes the top-response network. A stringent threshold (top-ranked 0.05 percentile of all paths) was used to identify highest perturbations. The networks were visualised in Cytoscape 3 (66). The functional categories of the genes present in the top-response network were assigned by mapping the functional category of its homologue in *M. tuberculosis* (67) from mycobrowser (https://mycobrowser.epfl.ch/)

#### Protein structure modelling and ligand docking

For protein ligand docking 3D x-ray crystal structure of PdtaS (PDB ID 2YKF (36)) was taken from RCSB PDB, a protein structure database (www.rcsb.org) (68). However the structure had a missing stretch of about 18aa from Val^91^ to Gly^108^ in the GAF domain. To model this missing stretch PDBeFold (69) was used to identify the structural fold similar to PdtaS and based on structure comparisons, 3DBA (70), a cGMP bound x-ray crystal structure of Gallus gallus was chosen. Using Modeller v9.18 (71) the missing portion of the PdtaS structure was reconstructed through multi template modelling using 3DBA and 2YKF. The PdtaS model thus built was energy minimised using GROMACS (72) with 2000 steps of steepest descent followed by 3000 steps of conjugate gradient algorithms. The quality of modelled structure was checked through ProSA (73) and further verified through a stereochemical quality check (74) and error estimation using a statistical scoring potential through DOPE (Discrete Optimized Protein Energy) and ERRAT score (75, 76). Three different pocket finding algorithms were used for identifying putative binding sites in the modelled protein, which are a) Fpocket (77) which identifies pockets through Voronoi tessellation, b) SITEHOUND (78) which is an energy based method for pocket identification and, c) pocketDepth (79) which is a grid based geometric method that identifies pockets through depth-based clustering. Cyclic di-GMP molecule (C2E) was docked onto the identified pockets using Autodock (80) and the pocket with the lowest binding energy was identified. Ligplot (81) was used to analyse protein-ligand interactions. To identify key residues that participate in ligand binding, ABS-scan (82) was used, which in essence mutates the residues in the binding pocket one by one to alanines, models the mutated proteins using Modeller followed by energy minimisation using GROMACS and C2E docking in the mutated binding pockets. Mutations which showed an aberration in ligand binding and a significant ΔΔG were selected for further experimental validation

#### In silico ligand screening

In order to identify the potential ligand candidate for modelled protein a computational screen was carried out. In brief the screen utilizes the 3D structural model of protein, identifies putative binding pockets in the protein, and scan it across all known small molecule ligand binding site in PDB. The screening was carried out using PocketMatch (83) which quantifies binding site similarity based upon structural descriptors such as shape and chemical nature of amino acid types at the site. Binding site is represented by 90 lists of sorted distances and aligned incrementally to obtain a similarity score called PocketMatch score (PMSmax) which range from 0 (no match) to 1 (perfect match). PMSmax of 0.6 and above is indicative of significant similarity between the binding sites. PMSmin score which reflects local structural similarity irrespective of relative size mismatch between two sites was used along with PMSmax for further analysis. Probable ligand hits were then compared structurally based upon tanimoto coeficient using Open Babel package, version 2.3.0 (84).

**Figure S1.**
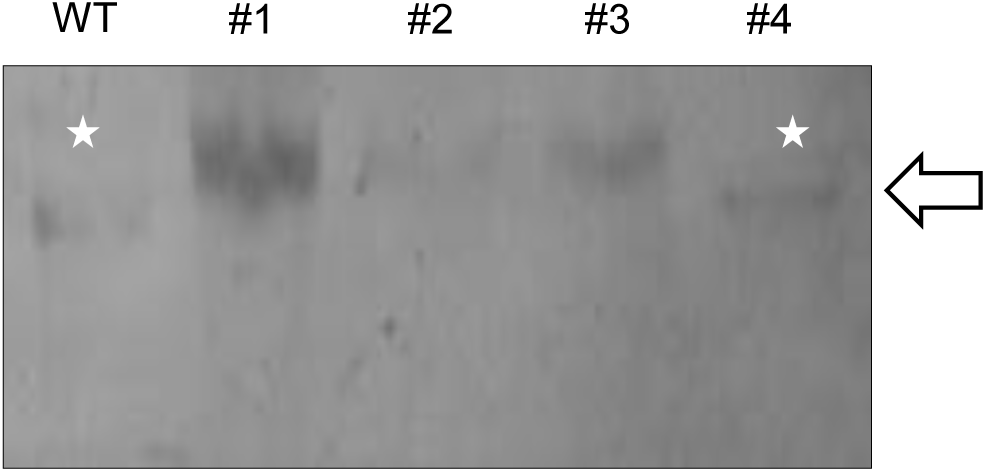
Confirmation of *pdtaS* knockouts using Southern Blotting. SacI digested genomic DNA isolated from wildtype and *pdtaS* knockout *M. smegmatis* colonies probed for replacement of *pdtaS* with a hygromycin – *sacB* cassette. White stars indicate bands expected from wildtype genomic DNA lacking the gene replacement.

**Figure S2.**
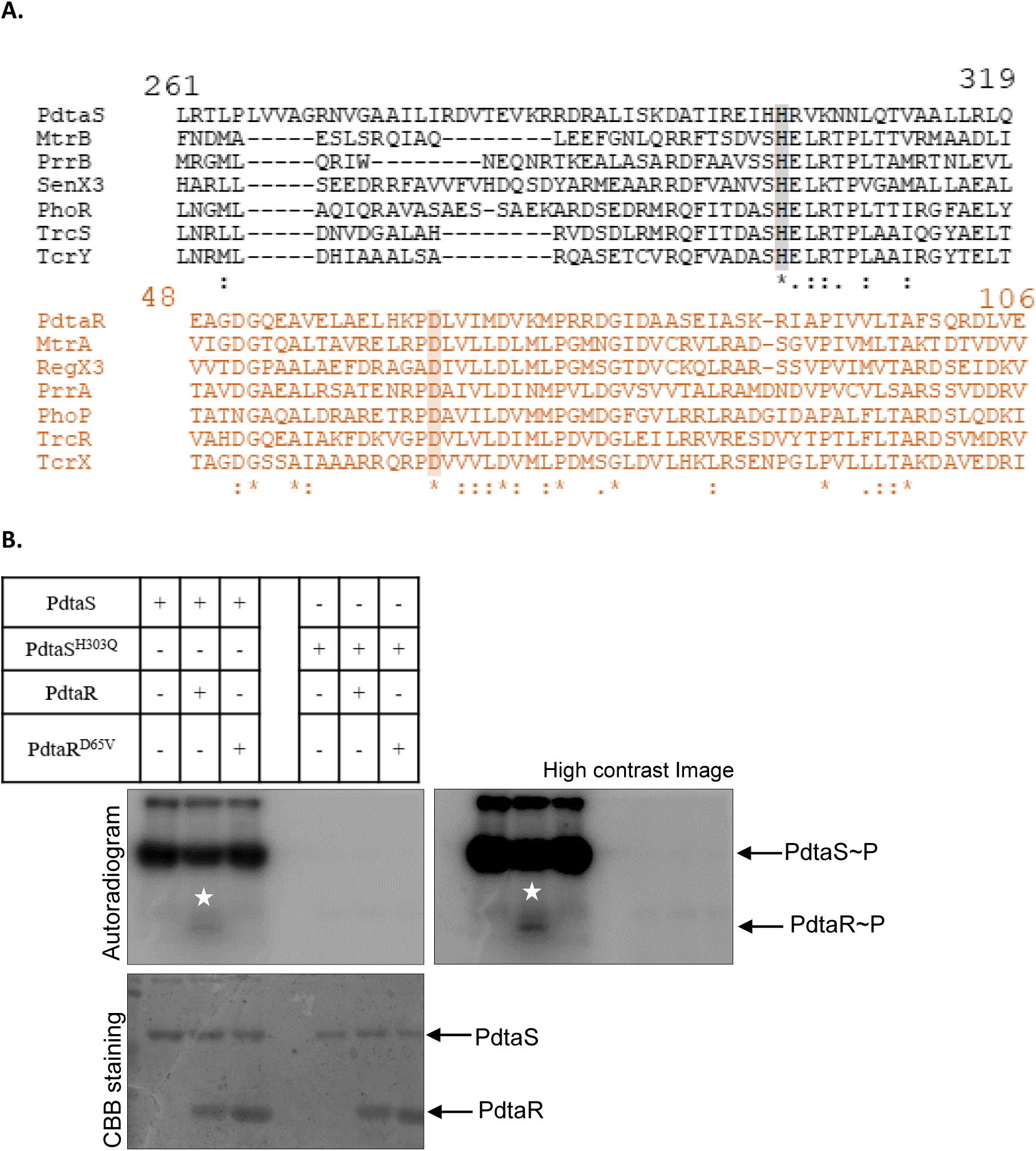
Identification and verification of phosphorylated residues. **A**. Multiple sequence alignment of PdtaS and PdtaR protein sequences with those of others *Mycobacterium tuberculosis* SKs and RRs respectively. Highlighted residues indicate the conserved amino acid involved in phosphorylation. **B**. Biochemical verification of phosphorylation sites by radiolabelled kinase assay. Wildtype PdtaS or PdtaS^H303^ was analysed for ability to undergo autophosphorylation and transfer phosphate to wildtype PdtaR or PdtaRD65V. Top panel represents the autoradiogram and bottom panel represents the corresponding Coomassie Brilliant Blue (CBB) staining. White stars indicate phosphorylated PdtaR bands.

**Figure S3.**
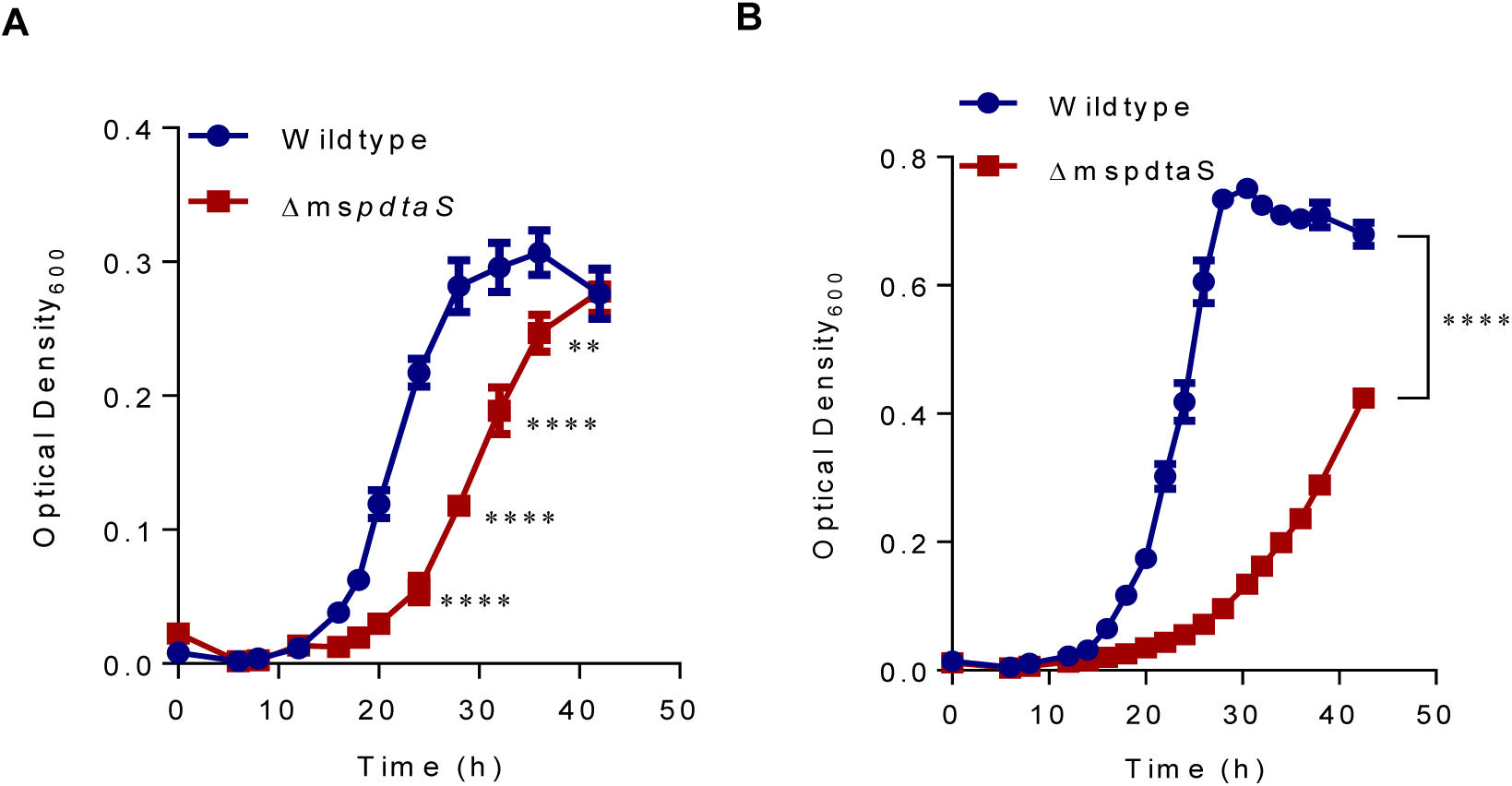
Growth defect of Δms*pdtaS* links to intracellular nutrient levels. The optical density at 600nm (O.D._600_) was measured at specific time points for Wildtype (WT), *pdtaS* knockout (Δms*pdtaS*) strains in a. 7H9 + glucose or b. 7H9 + glycerol after 8h starvation of pre-inoculum cultures grown in tryptic soy broth. Error bars represent the standard error of the mean of biological triplicates. Statistical significance as measured by unpaired t-tests for each row is depicted as * – p<0.01, **** – p<0.0001.

**Figure S4.**
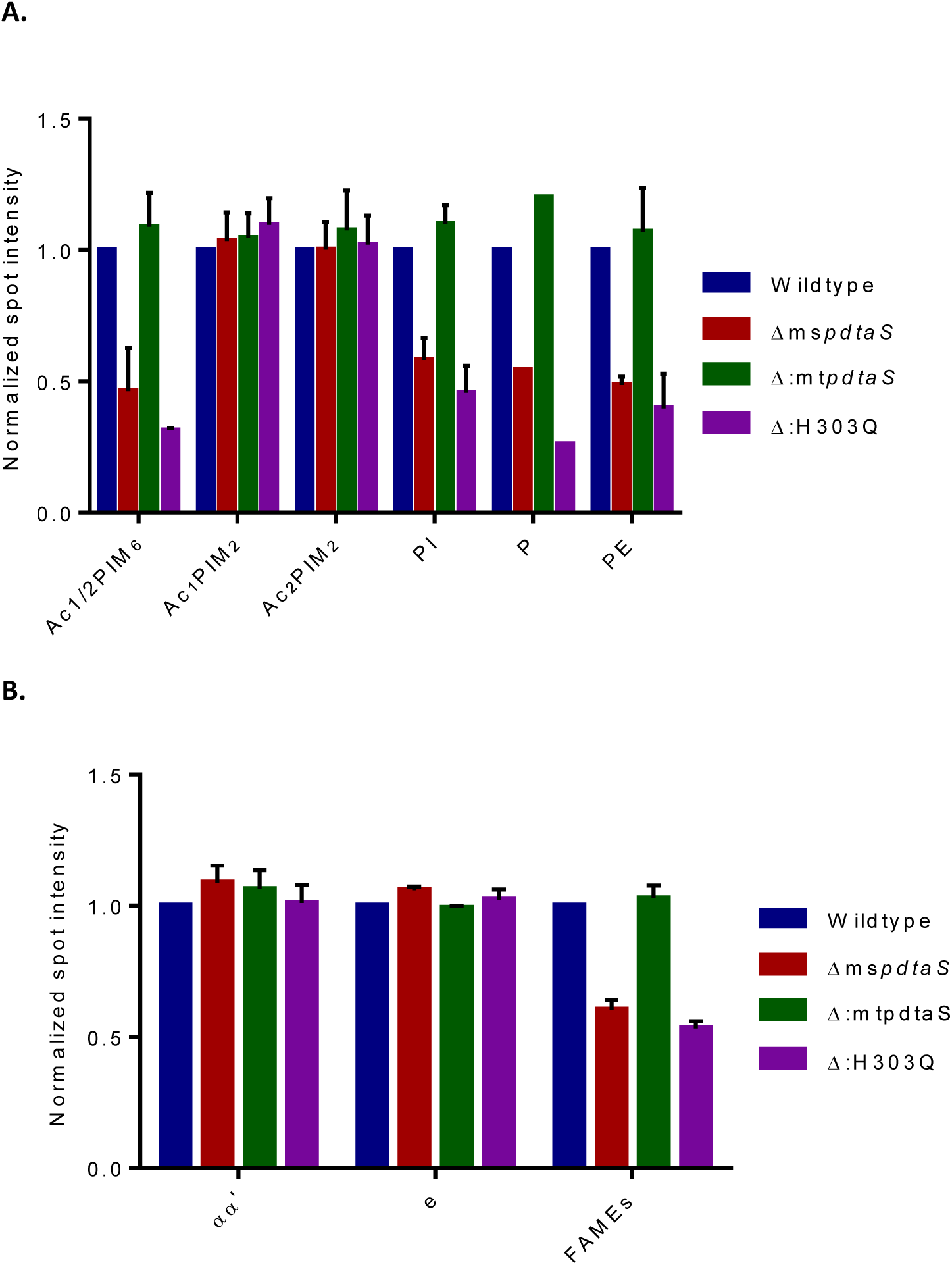
Normalized intensities of lipids. a. Spot intensities of individual lipids separated on 2D-TLCs using system E and b. Total FAMEs and MAMEs separated on 1D-TLCs. a. Intensities of Ac1PIM6 and Ac2PIM6 are clubbed together as Ac1/2PIM6 due to poor resolution in some experiments. Spot intensities of P are values from a single experiment only. All spots were normalized to wildtype. AcPIM– acylated phosphatidylinositol mannoside (integers denote the number of mannoside and acyl groups), PI – phosphatidyl inositol, P – phospholipid, PE – phosphatidyl ethanolamine. Error bars represent standard error of means across duplicates (n = 2). b. α and α’ mycolates were clubbed together as a single αα’ band due to lack of resolution between the bands. ε – epoxy. n = 3.

**Figure S5.**
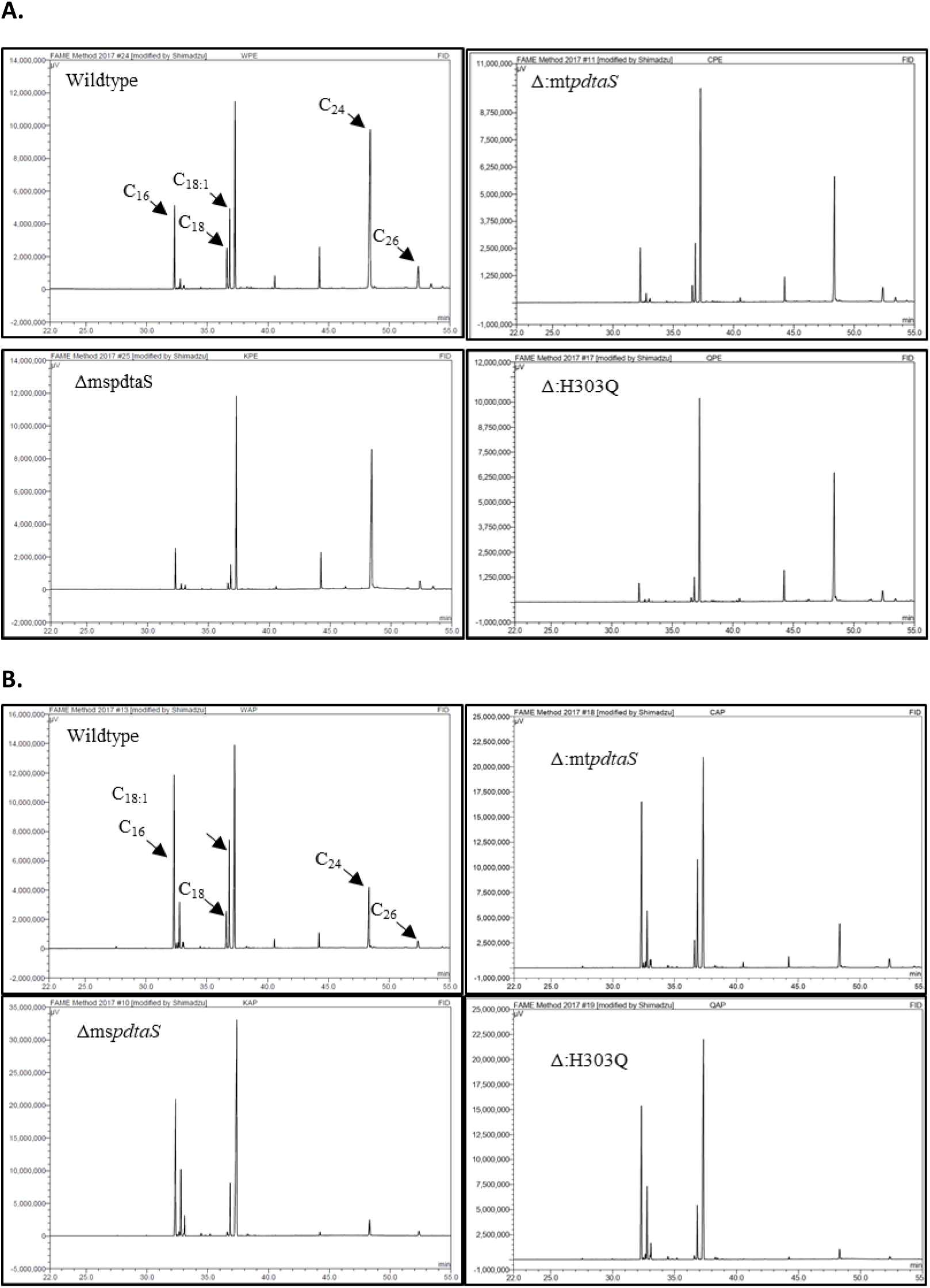
GC-MS analysis of apolar FAMEs. a.Intracellular apolar and b. extracellular apolar FAMEs from Wildtype, pdtaS knockout (ΔmspdtaS), complemented (Δ:mtpdtaS) and phosphorylation defective (Δ:H303Q) strains were studied using GC-MS. Peak annotation was calibrated using Sigma-Aldrich fatty acid methyl esters as standards (n = 1).

**Figure S6.**
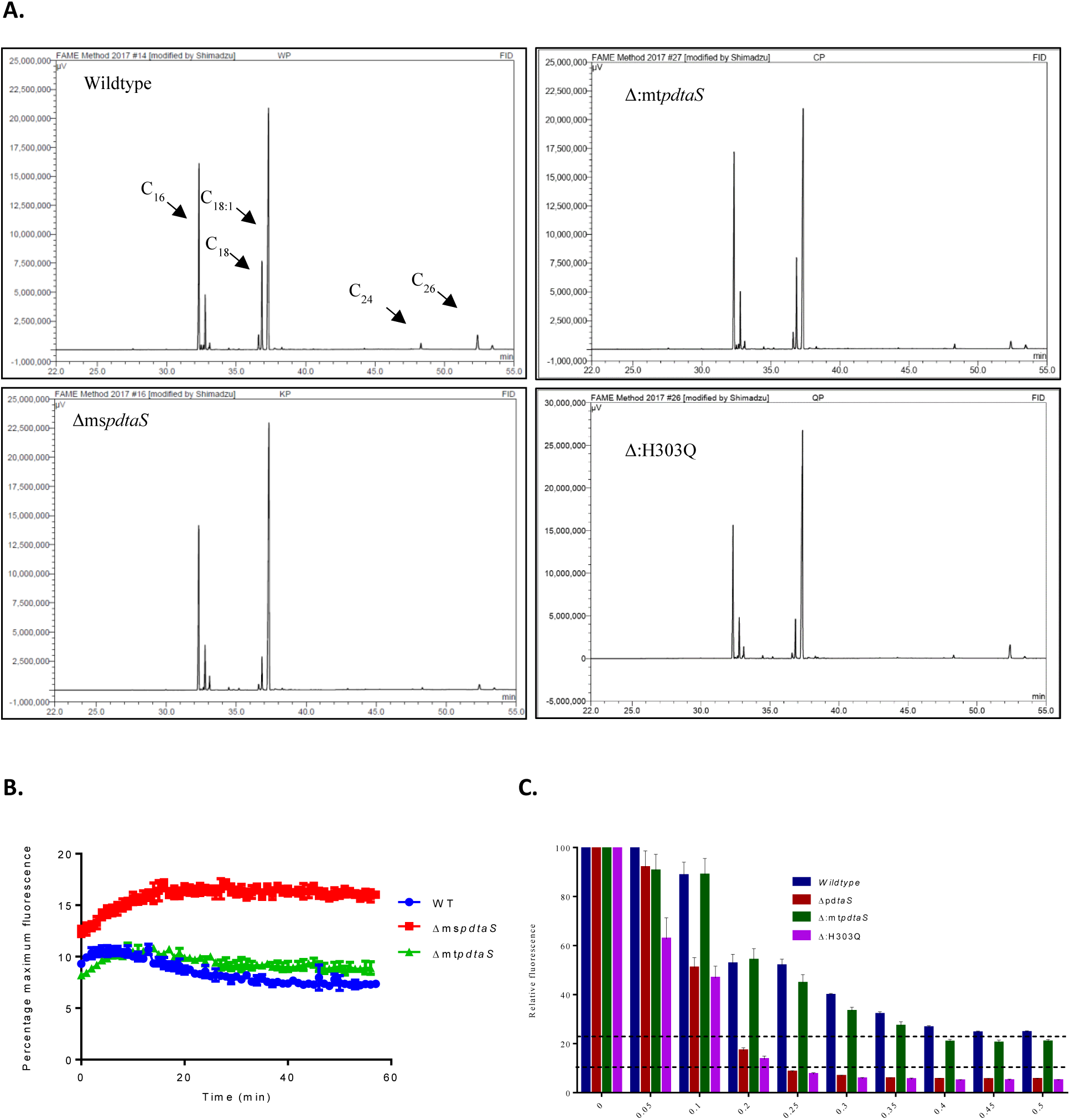
GC-MS analysis of polar intracellular FAMEs, Membrane permeability and antibiotic sensitivity. a. Polar FAMEs from Wildtype, pdtaS knockout (ΔmspdtaS), complemented (Δ:mtpdtaS) and phosphorylation defective (Δ:H303Q) strains were studied using GC-MS. Peak annotation was calibrated using Sigma-Aldrich fatty acid methyl esters as standards (n = 1). **b.** Measurements of membrane permeability through ethidium bromide uptake and **c.** sensitivity to streptomycin a. Fluorescence intensities of ethidium bromide (EtBr) (ex. 530nm em. 590nm) was measured over time for Wildtype (WT), *pdtaS* knockout (Δms*pdtaS*) and complemented (Δ:mt*pdtaS*) strains incubated with 1µg/ml EtBr. b. Wildtype (blue), mutant (red), Δ:mtpdtaS (green) and Δ:H303Q (purple) strains were grown in poor medium supplemented with cas amino acids in the presence of various concentrations of streptomycin in a 96-well plate. Black dashed lines indicate MIC values for mutant and wildtype cells. Error bars represent the standard error of the mean of biological triplicates (n = 3).

**Figure S7.**
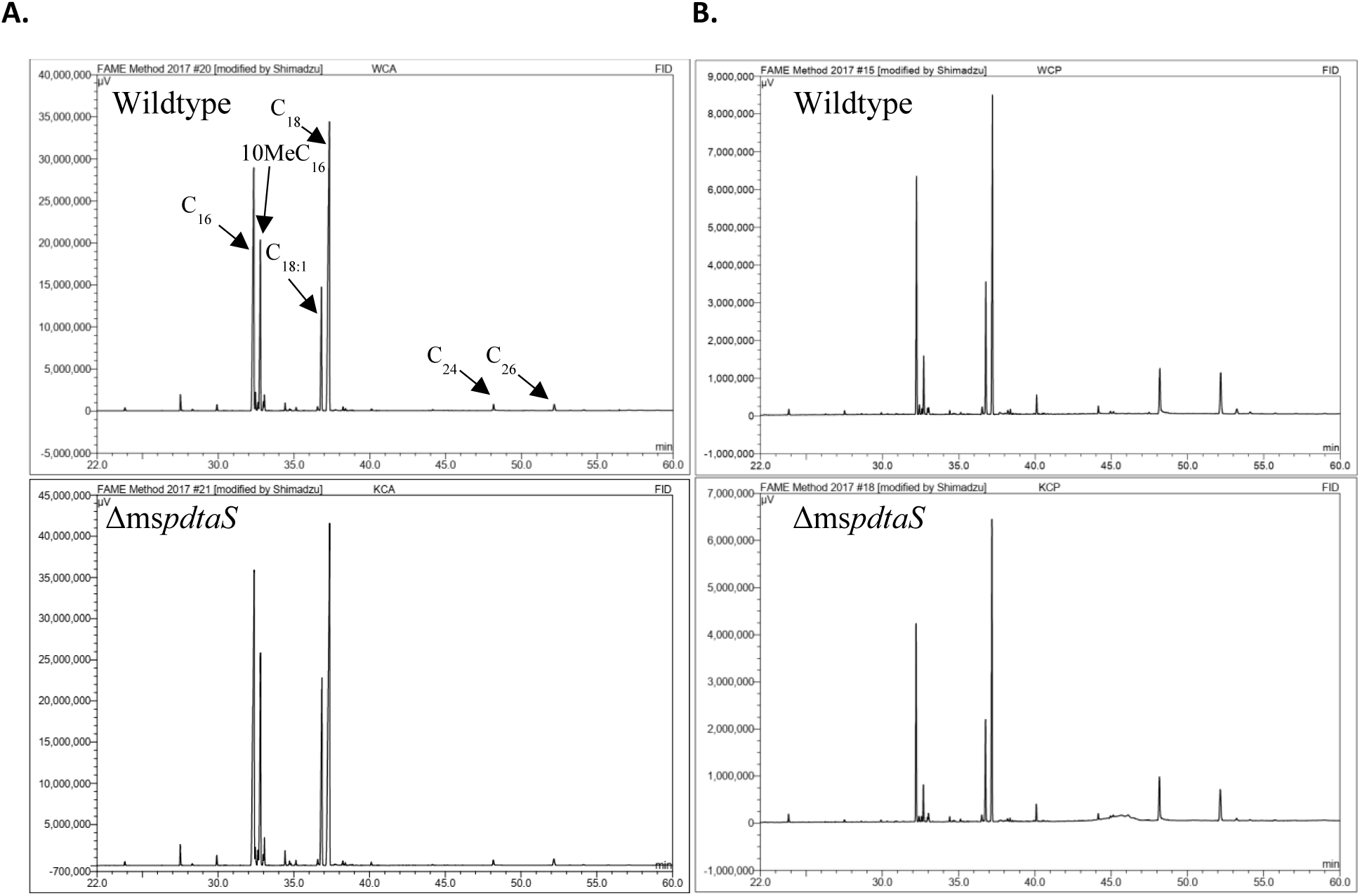
GC-MS analysis of apolar and polar FAMEs with cas supplementation. Total **a.** apolar and **b.** polar FAMEs from Wildtype and pdtaS knockout (ΔmspdtaS) strains grown with cas amino acid supplementation were studied using GC-MS. Peak annotation was calibrated using Sigma-Aldrich fatty acid methyl esters as standards (n = 1).

**Figure S8.**
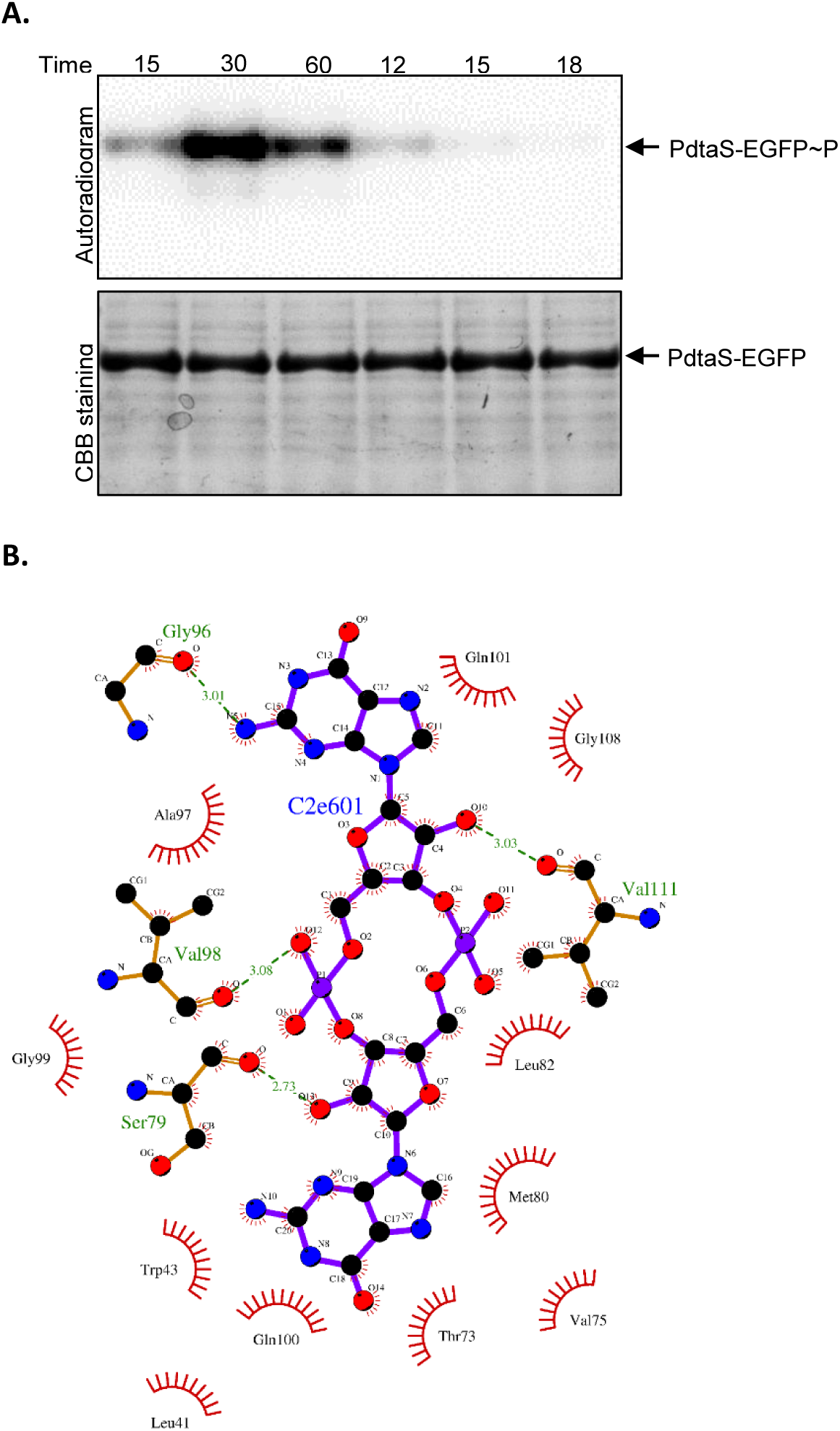
Kinetics of PdtaS-EGFP autophosphorylation and Interactions of c-di-GMP with the PdtaS binding pocket. **a.** Autophosphorylation time course analysis for PdtaS-EGFP. The top panel shows the autoradiogram and corresponding coomassie brilliant blue (CBB) staining. **b.** Detailed interaction map of ligand (Violet trace) and protein residues (Brown trace) in the binding pocket. Hydrogen bonds between interacting residues are represented as dashed lines, hydrophobic contacts are represented by arcs with radiating spokes.

**Figure S9.**
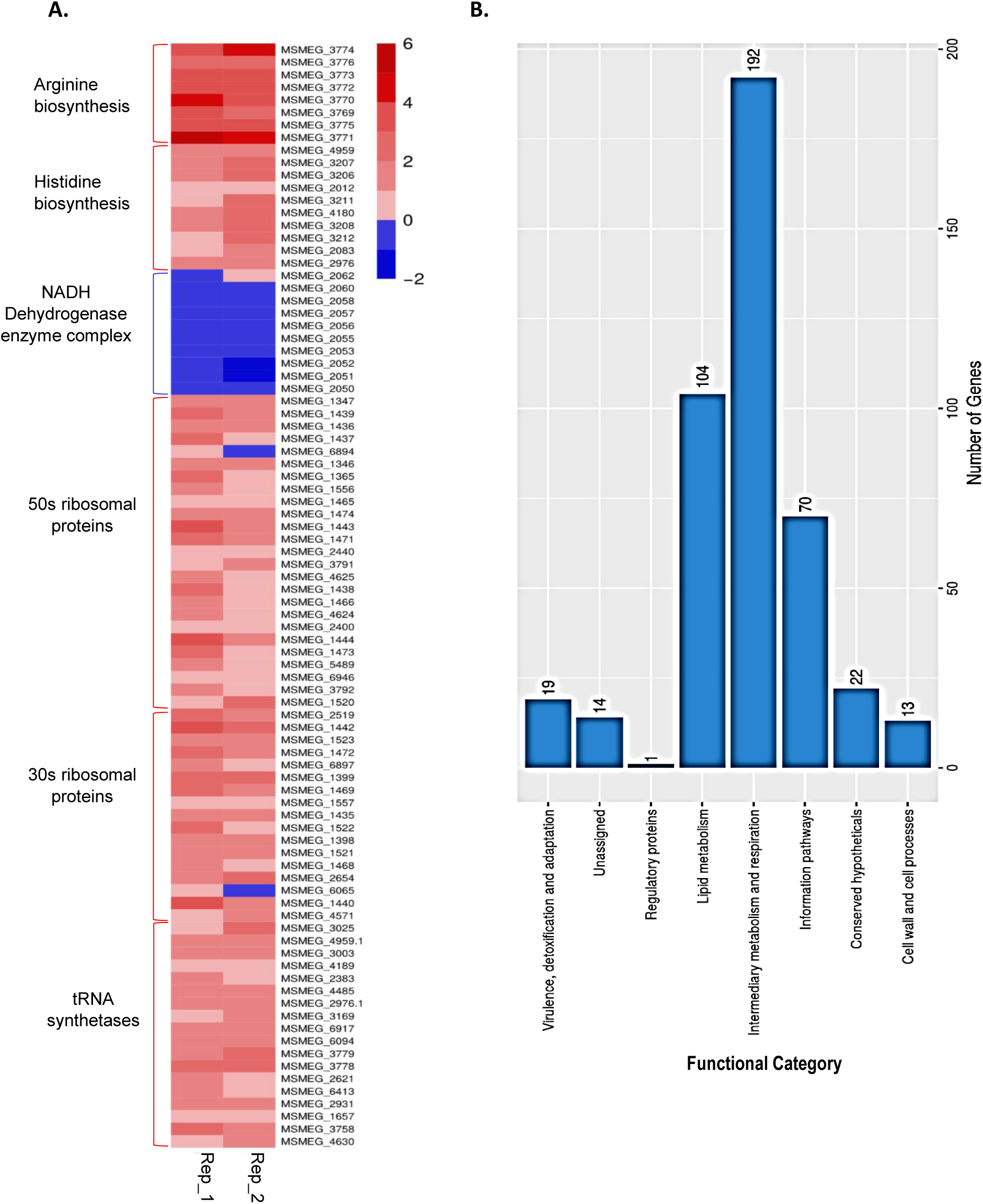
Heatmap of selected genes and frequency bar plot of gene functional categories in the top-response network. a. The fold change of arginine biosynthesis operon, histidine biosynthesis operon, NADH dehydrogenase enzyme complex operon, 50s ribosomal protein operon, 30s ribosomal protein operon and tRNA synthetases in ΔpdtaS cells when compared to wildtype M. smegmatis is shown. Colours correspond to degree of fold change from blue (downregulated) to red (upregulated). **b.** Genes in the top response network were sorted into functional categories to identify the major processes enriched in *pdtaS* knockout cells.

