## Supplementary File 2 for "The histidine kinase PdtaS is a cyclic di-GMP binding metabolic sensor that controls mycobacterial adaptation to nutrient deprivation"

**List of metabolites enriched in the *pdtaS* knockout**

| **Metabolites** | **Metabolite class** | **Fold change** |
| --- | --- | --- |
| DG (Diacylglycerol)[33:2] | Acyl glycerides | 3.49 |
| TG (Triacylglycerol)[62:0] | Acyl glycerides | 2.95 |
| TG[64:0] | Acyl glycerides | 2.18 |
| DG[35:1] | Acyl glycerides | 2.12 |
| TG[66:0] | Acyl glycerides | 2.11 |
| 3,4-Dihydroxymandelic acid | Aromatic metabolites | Absent in WT |
| 3-Phenylpropionylglycine | Aromatic metabolites | Absent in WT |
| CL (Cardiolipin)[57:4] | Cardiolipins | 5.09 |
| Linoleoylcholine | Choline metabolism | 2.32 |
| 16,18-Oxo-18-CoA-dinor-LTE4 | CoA metabolism | 3.36 |
| 18Hydroxy-20-oxo-20-CoA-LTE4 | CoA metabolism | 2.88 |
| 18-oxo-18-CoA-dinor-LTE4 | CoA metabolism | 2.07 |
| Tetradecadienoic acid | Fatty acids, alcohols and esters | 35.48 |
| trimethyl-hexadecatetraenoic acid" | Fatty acids, alcohols and esters | 3.02 |
| 3,12-dihydroxy palmitic acid | Fatty acids, alcohols and esters | 2.52 |
| Dodecanoic acid | Fatty acids, alcohols and esters | 2.19 |
| Hydroxy-tetradecanoic acid | Fatty acids, alcohols and esters | 2.00 |
| Ganglioside GM2 (d18:1/23:0) | Gangliosides | 9.68 |
| Ganglioside GM3 (d18:0/22:0) | Gangliosides | 6.48 |
| Ganglioside GM3 (d18:0/20:0) | Gangliosides | 5.20 |
| Ganglioside GM3 (d18:0/23:0) | Gangliosides | 5.17 |
| Ganglioside GM3 (d18:1/25:0) | Gangliosides | 4.88 |
| Ganglioside GA2 (d18:1/24:0) | Gangliosides | 4.40 |
| Ganglioside GM3 (d18:0/18:0) | Gangliosides | 3.91 |
| Ganglioside GM3 (d18:0/26:1) | Gangliosides | 3.46 |
| Ganglioside GM3 (d18:0/24:1) | Gangliosides | 3.21 |
| Ganglioside GM3 (d18:1/23:0) | Gangliosides | 3.16 |
| Ganglioside GM3 (d18:1/26:1)) | Gangliosides | 3.04 |
| Ganglioside GA1 (d18:1/25:0) | Gangliosides | 2.97 |
| Ganglioside GA2 (d18:1/26:1) | Gangliosides | 2.51 |
| Ganglioside GM3 (d18:0/25:0) | Gangliosides | 2.16 |
| PS (Phosphatidylserine)(O-16:0/14:0) | Glycerophospholipids | 2.09 |
| 1-Deoxy-D-xylulose 5-phosphate | Mixed class | Absent in WT |
| 7-Methylinosine | Mixed class | Absent in WT |
| Canavaninosuccinate | Mixed class | Absent in WT |
| Tetrahydropentoxyline | Mixed class | 11.23 |
| N-Acetyldopamine | Mixed class | 5.45 |
| PG (Phosphatidylglycerol)(P-16:0/0:0) | Mixed class | 3.59 |
| Isoleucyl-Phenylalanine | Mixed class | 2.90 |
| 25-hydroxycholecalciferol 6,19-sulfur dioxide adduct | Mixed class | 2.88 |
| PS[38:0] | Mixed class | 2.33 |
| PI (Phosphatidylinositol)(O-16:0/19:1) | Mixed class | 2.16 |
| 8-Hydroxy-7-methylguanine | Nucleosides and nucleotide metabolism | Absent in WT |
| 5'-Methylthioadenosine | Nucleosides and nucleotide metabolism | 10.52 |
| Cyclosporin A | Other class | 3.70 |
| DGDG[54:6] | Other class | 3.19 |
| Aspartyl-Methionine | Peptides | Absent in WT |
| Histidinyl-Methionine | Peptides | Absent in WT |
| Ornithino-L-alanine | Peptides | Absent in WT |
| Phenylalanyl-Arginine | Peptides | 2.68 |
| 1(3)-glyceryl-PGD2 | prostaglandins, leukotrienes and resolvins | 2.35 |
| 2-trans,6-cis,10-trans-geranylgeranyl diphosphate | Sterol and steroid metabolism | 3.48 |
| 1alpha-hydroxy-18-(4-hydroxy-4-methylpentyloxy)-23,24,25,26,27-pentanorcholecalciferol | Sterol and steroid metabolism | 2.16 |
| lithocholic acid sulfate | Sterol and steroid metabolism | 2.10 |
| (S1)-Methoxy-3-heptanethiol | Sulphur containing metabolites | Absent in WT |
| Acetylcysteine | Sulphur containing metabolites | Absent in WT |

**List of metabolites depleted in the *pdtaS* knockout**

| **Metabolites** | **Metabolite class** | **Fold change** |
| --- | --- | --- |
| Decenoylcarnitine | Acyl carnitine | 0.33 |
| TG[64:11] | Acyl glycerides | 0.46 |
| DG[39:3] | Acyl glycerides | 0.45 |
| TG[58:3] | Acyl glycerides | 0.45 |
| DG[43:1] | Acyl glycerides | 0.40 |
| TG[64:17] | Acyl glycerides | 0.40 |
| DG[44:4] | Acyl glycerides | 0.31 |
| TG[55:6] | Acyl glycerides | 0.30 |
| TG[64:15] | Acyl glycerides | 0.29 |
| TG[62:16] | Acyl glycerides | 0.26 |
| TG[60:3] | Acyl glycerides | 0.24 |
| TG[59:4] | Acyl glycerides | 0.20 |
| TG[66:17] | Acyl glycerides | 0.14 |
| TG[57:4] | Acyl glycerides | 0.11 |
| Cer (Cerramide)(d18:1/23:0) | Ceramides and sphingolipids | 0.38 |
| Behenic acid | Fatty acids, alcohols and esters | 0.48 |
| 1,2-eicosanediol | Fatty acids, alcohols and esters | 0.46 |
| Tetradecanedioic acid | Fatty acids, alcohols and esters | 0.45 |
| Eicosatrienoic acid isobutylamide | Fatty acids, alcohols and esters | 0.45 |
| 1,2-heneicosanediol | Fatty acids, alcohols and esters | 0.37 |
| Oxododecanoic acid | Fatty acids, alcohols and esters | 0.34 |
| methyl-octadecadienoic acid | Fatty acids, alcohols and esters | 0.28 |
| dimethyl-hexadecadienoic acid | Fatty acids, alcohols and esters | 0.24 |
| Lauryl linolenate | Fatty acids, alcohols and esters | 0.11 |
| Ganglioside GM3 (d18:0/24:0) | Gangliosides | 0.49 |
| PS(O-20:0/22:0) | Glycerophospholipids | 0.48 |
| PG(O-20:0/20:3) | Glycerophospholipids | 0.48 |
| LysoPC[16:0] | Glycerophospholipids | 0.47 |
| PC (Phosphatidylcholine)(o-18:0/24:0) | Glycerophospholipids | 0.37 |
| PS[44:4] | Glycerophospholipids | 0.35 |
| PA[44:2] | Glycerophospholipids | 0.32 |
| PG(O-16:0/22:2) | Glycerophospholipids | 0.26 |
| PS-NAc[52:1] | Glycerophospholipids | 0.21 |
| PC[43:4] | Glycerophospholipids | 0.18 |
| PA (Phosphatidic acid)[44:0] | Glycerophospholipids | 0.17 |
| PS(O-20:0/22:2) | Glycerophospholipids | 0.16 |
| 1alpha,25-dihydroxy-22,23,24,24a-tetradehydro-24a-homocholecalciferol | Mixed class | 0.47 |
| 8-Hydroxygeraniol 8-O-glucoside | Mixed class | 0.46 |
| PG(P-20:0/22:2) | Mixed class | 0.44 |
| Verazine | Mixed class | 0.40 |
| PG(P-20:0/22:2) | Mixed class | 0.40 |
| Galabiosylceramide (d18:1/24:1) | Mixed class | 0.38 |
| PE (Phosphatidylethanolamine)[46:7] | Mixed class | 0.34 |
| PI-Cer(t20:0/26:0(2OH)) | Mixed class | 0.31 |
| Dodecaprenyl phosphate-galacturonic acid | Mixed class | 0.31 |
| PS(P-20:0/22:4) | Mixed class | 0.29 |
| Galabiosylceramide (d18:1/26:1) | Mixed class | 0.29 |
| PG[32:2] | Mixed class | 0.24 |
| PG(O-18:0/22:2) | Mixed class | 0.15 |
| PG[32:1] | Mixed class | 0.14 |
| MIPC(t18:0/22:0(2OH)) | Mixed class | 0.13 |
| LBPA (Lysobisphosphatidic acid)[34:2] | Mixed class | 0.10 |
| (9Me,-d19:3)sphingosine | Mixed class | 0.07 |
| PI(P-20:0/19:1) | Mixed class | 0.07 |
| PG(O-20:0/22:2) | Mixed class | 0.05 |
| 2-(3-Carboxy-3-(methylammonio)propyl)-L-histidine | Other class | 0.47 |
| Spermidine | Other class | 0.37 |
| Theonellasterol B | Other class | 0.36 |
| LacCer(d18:0/26:0) | Other class | 0.35 |
| 2,2,6,6-Tetramethyl-4-piperidinone | Other class | 0.30 |
| DAT(16:0/24:0(2Me[S],3OH[S],4Me[S],6Me[S])) | Other class | 0.20 |
| PI[44:1] | Phosphoinositol lipids | 0.41 |
| PI[43:1] | Phosphoinositol lipids | 0.21 |
| Dehydroergosterol | Sterol and steroid metabolism | 0.45 |
| 8,18-propano-retinal | Sterol and steroid metabolism | 0.45 |
| 1alpha,19,25-trihydroxy-10,19-dihydrocholecalciferol | Sterol and steroid metabolism | 0.39 |
| 1alpha-hydroxy-24-oxo-26,27-cyclocholecalciferol | Sterol and steroid metabolism | 0.21 |
